# Distinct distributions of myosin motor conformations during contraction of slow and fast skeletal muscle

**DOI:** 10.1101/2025.10.13.679501

**Authors:** Cameron Hill, Michaeljohn Kalakoutis, Alice Arcidiacono, Yanhong Wang, Elisabetta Brunello, Luca Fusi, Malcolm Irving

## Abstract

Slow skeletal muscles maintain posture and produce graded movement at low metabolic cost. Force development and ATP utilisation during fixed-end contractions are typically five times slower in slow than fast muscles from the same species. Mechanical measurements previously suggested that more myosins are attached to thin filaments during contraction of slow muscle, which seems incompatible with its high efficiency. We therefore used small-angle X-ray diffraction to provide a structural estimate of the fraction of myosins attached to thin filaments in slow muscle. X-ray signals associated with myosin binding to actin indicate that only about 10% of myosin motors are actin-bound during fixed-end tetani of rat soleus slow muscles, compared with about 25% in mouse EDL fast muscle. Moreover, X-ray signals associated with the helical organisation of OFF myosin motors in the thick filaments show that about 70% of myosin motors remain in the OFF conformation during tetanic contraction of slow muscle, compared with only 30% in fast muscle. The much slower force development in soleus muscle also allowed clear separation of early structural changes in thick filaments on activation, some of which are distinct from those reported previously in fast muscles. Moreover, the early structural changes in soleus muscle have about the same amplitude in a twitch and a tetanus, suggesting that they are triggered by thin filament activation rather than thick filament stress, and implying a fast signalling pathway between thin and thick filaments.

**Key Points:**

- The interaction between myosin motors and actin filaments in slow skeletal muscles maintain posture and produce graded movement at low metabolic cost.
- Mechanical studies have suggested that more myosins are attached to actin filaments in slow than in fast muscle, but this seems incompatible with its high efficiency.
- We used X-ray diffraction to show that there are fewer myosin motors attached to actin in slow muscle than in fast muscle because more motors are sequestered on the myosin filament.
- The slower force development in slow muscle also allowed us to isolate and characterise fast changes in myosin motor conformation associated with activation of the actin filaments.
- The results reveal a distinct pathway of inter-filament signalling in slow muscle that could help the development of novel therapies for muscle weakness.

## Introduction

Slow skeletal muscles maintain steady tension and produce graded motion at low metabolic cost, in contrast with the more ballistic and metabolically expensive movements driven by fast skeletal muscles. Slow muscles shorten more slowly against an external load and develop force more slowly in fixed-end or isometric contractions. In the most extensively studied slow and fast muscles, the soleus and extensor digitorum longus (EDL) muscles of the mouse, the rates of isometric force development and ATP utilisation differ by a factor of five (Barclay *et al*., 1993). The same roughly five-fold difference in the rate of isometric ATP utilisation is observed in single demembranated soleus and EDL fibres of the rat during maximal calcium activation (Bottinelli *et al*., 1994).

Slow or type-1 muscle fibres like those in soleus express predominantly the MYH7 myosin heavy chain, and the fast type 2A or 2X fibres in EDL express MYH2 and MYH1 respectively (Weiss *et al*., 1999; Dos Santos *et al*., 2022). Isolated type-1 myosin hydrolyses ATP more slowly in the presence of actin than type-2 myosin, and the rate constants for ATP hydrolysis on myosin and for the release of ADP from the actin-myosin complex are five to ten times smaller (Iorga *et al*., 2007). Although fast and slow muscles also express different isoforms of proteins involved in calcium signalling and thin filament regulation, calcium release in response to single action potential stimulation is more than sufficient to saturate the regulatory sites on troponin in both muscle types (Carroll *et al*., 1997; Baylor & Hollingworth, 2003). Moreover, intracellular free calcium concentration [Ca^2+^]_i_ peaks only 2-3 ms after stimulation in both soleus and EDL at 28°C, much faster than isometric force generation in the soleus fibres, which has a half-time of about 20 ms in those conditions. Differences in calcium signalling and thin filament regulation seem to make little contribution to the different rates of force generation and ATP utilisation in fast and slow muscles.

The rate and extent of force generation in fast skeletal muscles also depends on activation of the myosin-containing thick filaments (Linari *et al*., 2015; Irving, 2017; Craig & Padrón, 2022; Brunello & Fusi, 2024). The myosin motor or head domains that generate force and shortening in their ATP-driven cyclic interaction with actin in the thin filaments are prevented from binding actin in resting muscle because they are folded back against their tails in a helical array on the surface of the thick filaments. In this sequestered or OFF state the ATPase activity of myosin is largely suppressed, minimising the metabolic cost of resting muscle (Stewart *et al*., 2010). In fast muscle fibres, myosin motors can be released from the OFF state by thick filament stress, leading to a positive feedback loop in which the first motors to become activated increase filament stress, releasing more motors which generate more stress and so on. This positive co-operativity has a clear functional advantage in the all-or-none ballistic action of fast skeletal muscles, but seems poorly adapted for the graded action of slow muscles, or for minimising the metabolic cost of slow muscle contraction. Those features, together with the absence of major fibre-type differences in calcium release and thin filament regulation, suggest a very different role for thick filament regulation in slow muscles: to limit the number of myosins that interact with actin and the associated ATP hydrolysis to the minimum required for tension maintenance and slow movement. One aim of the present work was to test that hypothesis using small-angle X-ray diffraction to determine the activation state of the thick filaments and the fraction of myosin motors attached to actin during isometric contraction of slow muscle.

Slow skeletal muscles also generate lower isometric force per cross-sectional area than fast muscles, typically by about a factor of two. For example, the isometric force produced during maximal calcium activation of demembranated fibres from rabbit soleus muscle at 25°C was reported as 165 kPa, compared with 317 kPa in fast fibres from rabbit psoas muscle (Percario *et al*., 2018; Caremani *et al*., 2022). Two very different explanations for the lower isometric force produced by slow muscle may be considered. One possibility is that individual fast and slow myosins produce the same unitary force (Palmiter *et al*., 1999), estimated as about 5 pN in recent studies (Woody *et al*., 2019; Shchepkin *et al*., 2020), but fewer myosin motors are attached to actin during fixed-end contractions of slow muscle. An alternative explanation arose from the observation that slow muscle fibres are less stiff than fast muscle fibres in rigor, i.e. in the absence of ATP, when all the myosins in the thick filament are assumed to be strongly bound to actin (Köhler *et al*., 2002; Brenner *et al*., 2012; Percario *et al*., 2018). If that assumption is correct, individual molecules of slow muscle myosin must be intrinsically less stiff than their fast muscle counterparts. It then follows from the stiffness of actively contracting fibres that the unitary force generated by slow muscle myosin would only be about 2 pN, and that a *greater* fraction of myosins, about 50% of the total present compared with about 30% in fast muscle, would be attached to thin filaments during fixed-end contraction of slow muscle (Percario *et al*., 2018). Those conclusions seem difficult to reconcile with the lower ATPase rate and high mechanical efficiency of slow muscle (Barclay *et al*., 2010*a*).

The experiments reported below aimed to distinguish between these alternatives and to determine the role of thick filament regulation in intact, electrically stimulated slow skeletal muscles. To do so we used time-resolved small-angle X-ray diffraction, which provides direct structure-based estimates of the degree of activation of the thick filaments and the fraction of myosins attached to thin filaments on the physiological timescale. We chose rat soleus muscles, which have a high proportion of slow type-1 fibres, for these studies, and compared the results with our previous X-ray studies of predominantly fast or type-2 EDL muscles of the mouse (Hill *et al*., 2021, 2022, 2025). We show that the level of thick filament activation in fixed-end tetanic contractions of the slow muscle is much less than in the fast muscle, and that fewer myosins are attached to thin filaments. Our results provide strong support for the hypothesis that the lower force and ATP utilisation of slow muscles are predominantly due to increased sequestration of myosins in the OFF state rather than a reduced force per myosin.

## Methods

### Animals and Muscle Preparation

Male rats (strain Wistar Han) were obtained from Charles River Laboratories (Wilmington, MA, USA) and housed at the European Synchrotron Radiation Facility (ESRF) Biomedical Facility, Grenoble, France, or the United Kingdom Health Security Agency (UKHSA), Didcot, UK, in 12:12 hour light:dark cycles at 20°C and 50% relative humidity, with *ad libitum* access to water and a standard lab diet. The rats used at ESRF were slightly older (9-11 weeks) than those at UKHSA (6-7 weeks).

Animals were sacrificed via cervical dislocation, followed by a confirmation method of permanent cessation of circulation by severing the femoral artery, in compliance with the UK Home Office Animals (Scientific Procedures) Act 1986, Schedule 1 and European Union regulation (directive 2010/63). After sacrifice, whole soleus muscles were dissected from the hindlimb under a stereomicroscope in a Sylgard dish continuously perfused with Krebs-Henseleit solution (composition in mM: NaCl 118; KCl 4.96; MgSO_4_ 1.18; NaHCO_3_ 25; KH_2_PO_4_ 1.17; glucose 11.1; CaCl_2_ 2.52) with a pH ∼7.4 at room temperature after equilibration with carbogen (95% O_2_, 5% CO_2_). Metal hooks were tied with 4-0 silk sutures at the proximal and distal tendons of the muscle to allow attachment to the experimental setup. The muscle was mounted in a custom 3D-printed resin chamber between a fixed hook and the lever of a dual-mode force/length transducer (300C-LR, Aurora Scientific, Aurora, Canada) and continuously perfused with Krebs-Henseleit solution equilibrated with carbogen at 27-28°C (Hill *et al*., 2025).

Electrical stimuli were provided by a high-power biphasic stimulator (701C, Aurora Scientific) via parallel platinum electrodes. The muscle was placed between a fixed mylar window and a second window attached to a 3D-printed screw to allow it to be positioned as close as possible to the muscle to minimise the X-ray path in the solution. The stimulus voltage was 1.5 times the required amount to elicit the maximum twitch force response. Optimal muscle length (*L*_0_) was set to produce maximum force in response to a 200 ms train of stimuli at 80 Hz repeated at 5-minute intervals. *L*_0_ was 24.6 ± 0.9 mm for the experiments at ESRF and 22.2 ± 0.6 mm for the experiments at Diamond (mean ± SD). Fibre length was assumed to be 69% of muscle length (Eng *et al*., 2008). Muscle cross-sectional area was estimated as *W*_MW_/(ρ\**L_0_*\*0.69), where ρ = 1.06 g.cm^-3^ is the density of the muscle, and *W*_MW_ is the muscle wet weight. For experiments at ESRF, *W*_MW_ was 111.3 ± 6.8 mg, giving a cross-sectional area of 6.2 ± 0.5 mm^2^. In experiments at Diamond, *W*_MW_ was 81.0 ± 2.9 mg giving a cross-sectional area of 5.2 ± 0.4 mm^2^. *L*_0_, *W*_MW_ and cross-sectional area were significantly larger for muscles used at ESRF (*t*-tests; P<0.05 for all). Peak force in fixed-end tetani at *L*_0_ at Diamond before X-ray exposure was 130.1 ± 17.4 kPa (mean ± SD; n = 4). Peak tetanic force could not be measured in the batch of muscles used at ESRF because it exceeded the range of the force transducer.

### Small-angle X-ray Diffraction Data Collection

The trough was sealed with silicon grease to prevent solution leakage, and the muscle was mounted vertically at *L*_0_ at beamline ID02 of the European Synchrotron Radiation Facility (ESRF, Grenoble, France) or beamline I22 of the Diamond Light Source (DLS, Didcot, Oxfordshire, United Kingdom) to take advantage of the smaller vertical beam focus to optimise spatial resolution along the meridional axis. The beamline and detector properties are provided in Table S1.

For the fixed-end twitch experiments, data were collected at beamline ID02, ESRF on a Eiger 2-4M detector (Dectris, Baden, Switzerland). The sample-to-detector distance was initially set to 31 m for alignment and to measure sarcomere length during the twitch, using 5 ms exposures and a 50 µm rhodium attenuator with 3% transmission. Rapid assessment of two-dimensional X-ray patterns was provided by SAXSutilities2 (Sztucki, 2021). Following alignment, the attenuators were replaced with a 50 μm molybdenum attenuator with 9% transmission for time-resolved experiments at 3.2 m.

For the fixed-end tetanus experiments, data were collected at the I22 beamline at DLS on a Pilatus P3-2M detector (Dectris). The sample-to-detector distance was set to 8.26 m throughout. Muscles were aligned at I22 with a 100 μm molybdenum attenuator with 0.5% transmission, and the assessment of patterns was provided by the Data Analysis WorkbeNch software (DAWN; Basham *et al*., 2015). The attenuator was then removed to provide an unattenuated beam for the time-resolved fixed-end tetanus experiments.

The detector position in each case was optimised so that X-ray reflections of interest did not fall in the gaps between detector tiles. The 8.26 m and 3.2 m sample-to-detector distances were used to measure changes in the equatorial reflections (11 and 10), the third-order meridional reflection (M3), the sixth-order meridional reflection (M6), the mixed first-order actin layer line (AL1) and myosin layer line (ML1), the sixth-order actin layer line (AL6), and layer-line sampling from the simple filament lattice in rat soleus muscle (Ma *et al*., 2019).

The stimulation protocol and force response are shown in Fig. 1A. Muscle length was set to *L*_0,_ and the muscle was electrically stimulated under fixed-end conditions for 237ms at 80 Hz to evoke a tetanus (Fig. 1A, black trace) or with a single stimulus to evoke a twitch (Fig. 1A, grey trace). Small-angle X-ray diffraction data were acquired in 64 frames for the tetanus, each with 8 ms integration and 2 ms latency time, or in 70 frames for the twitch, each with 4.5 ms integration and 0.5 ms latency. To minimise radiation damage, X-ray exposure was limited by a fast shutter at both beamlines and the muscle moved vertically and horizontally between successive X-ray exposures. X-ray data were added from 40 to 60 contractions per muscle for twitch experiments, and 2 to 7 contractions per muscle for tetanus experiments. Records in which force had declined by more than 15% from the first record, the quality of the diffraction pattern substantially deteriorated relative to the first record, or in which collagen-based reflections were seen, indicating the presence of tendon in the X-ray beam, were excluded from further analysis.

**Figure 1.**
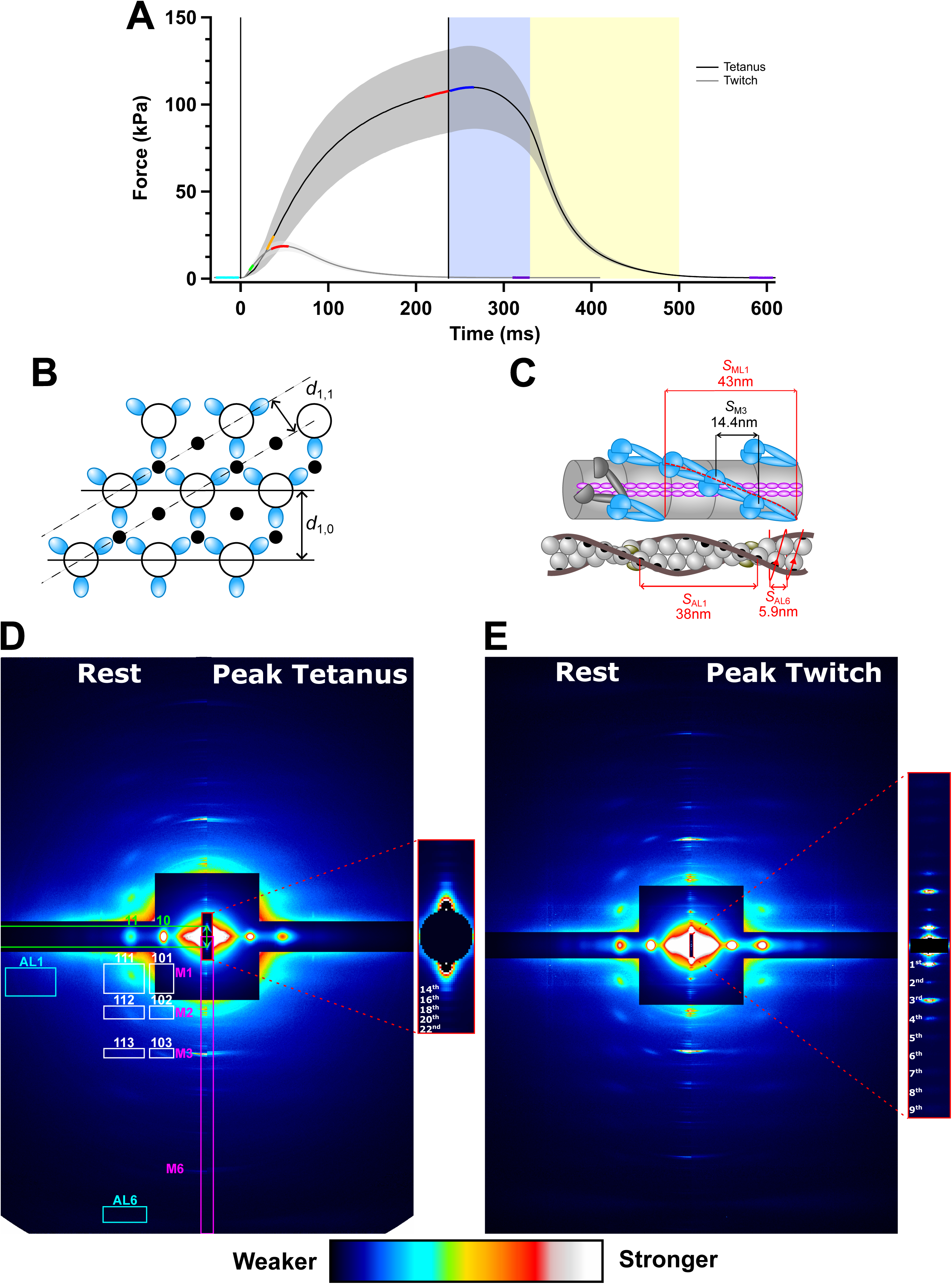
Mechanical protocol, thick and thin filament structure, and small-angle X-ray diffraction patterns from intact rat soleus muscle. *A*, Force per cross-sectional area in response to a single electrical impulse (twitch, grey trace) and a 237 ms train of electrical stimuli under fixed-end conditions (tetanus, black). The blue shaded region denotes isometric relaxation, and the yellow shaded region the subsequent transition to exponential relaxation. Coloured segments of the force trace denote periods used for averaging the X-ray data: cyan, rest; orange, early activation during the tetanus; green, early activation during the twitch; red, peak force; blue, isometric relaxation in the tetanus; purple, mechanically relaxed. Mean ± SD for n = 4 muscles in the tetanus and n = 7 in the twitch. *B*, Transverse section showing the hexagonal lattice of thick (open circles) and thin (filled circles) filaments and the lattice planes associated with the equatorial-based 10 and 11 reflections. Blue ellipses represent the three pairs of myosin motors which have the same azimuth at each axial level of the thick filaments, giving rise to discrete off-axis reflections. *C*, longitudinal view of the myosin-containing thick filament (upper) and actin-containing thin filament (lower) in resting muscle. Myosin motor dimers (blue) are folded against their tails in a three-stranded helical array on the surface of the thick filament backbone (grey cylinder). The axial periodicity of the helix is ca 43 nm; that of the motors is ca 14.4 nm. Some motors in resting muscle (grey ellipses) do not join the helical array. Actin monomers (grey spheres) are arranged in a double-stranded, long-pitch helix with an axial periodicity of about 38 nm which can also be described as a right-handed genetic helix with a periodicity of about 5.9 nm. Tropomyosin (brown) inhibits myosin binding by covering the binding sites on actin for myosin (black circles). *D*, Small-angle X-ray diffraction pattern of rat soleus muscle at rest (left) and at peak force in the tetanus (right) recorded at beamline I22 of the Diamond Light Source with a sample-to-detector distance of 8.2 m, mirrored horizontally only, from the average of three frames at rest and peak force from 20 records for 4 muscles, equivalent full-beam exposure, 480 ms. Magenta box: integration region for the first- to sixth-order myosin-based meridional reflections M1, M2, M3 and M6. Green box: integration region for the equatorial reflections 10 and 11. Cyan boxes: integration regions for first- and sixth-order actin-based layer line reflections AL1 and AL6. White boxes: Integration regions for off-axis myosin-based reflections. Red box: Integration region for the sarcomere reflections. *Inset,* ultra-small-angle X-ray diffraction region for resting muscle, showing 14^th^ to 22^nd^ even-order sarcomere reflections. *E*, Two-dimensional small-angle X-ray diffraction pattern of rat soleus at rest (left) and at peak force in the twitch (right) collected at beamline ID02 of the ESRF, with a sample-to-detector distance of 3.2 m, mirrored horizontally and vertically, average of four frames at rest and peak force from 145 records in four muscles; 234.9 ms. *Inset,* Ultra-small-angle X-ray diffraction patterns recorded from resting muscle with a sample-to-detector distance of 31 m, from 15 records in seven muscles, full beam equivalent exposure 8.1 ms. Sarcomere orders one to nine are labelled.

Force, stimulus, muscle length, and X-ray acquisition timing were sampled and analysed using custom-made software written in LabVIEW (National Instruments).

### X-ray Diffraction Data Analysis

Small-angle X-ray diffraction patterns were analysed using DAWN, SAXSutilities2, SAXS package (P. Boesecke, ESRF, Grenoble, France), Fit2D (A. Hammersley, ESRF, Grenoble, France), ImageJ (National Institute of Health, Bethesda, USA; Schneider *et al*., 2012) and Igor Pro 8 (WaveMetrics Inc., Portland, USA).

### Analysis of Ultra-Small-Angle X-Ray Diffraction Patterns

For experiments at ID02, the sarcomeric X-ray reflections were recorded with a sample-to-detector distance of 31 m at three places along the long axis of each muscle. Following subtraction of the camera background, diffraction data were integrated 0.56 µm^-1^ on either side of the meridional axis, and the residual background intensity was removed using an arc hull algorithm without smoothing using the Baselines extension in Igor Pro. The axial region 0.25-0.57 µm^-1^ containing the first-order sarcomere reflection was fitted with a single Gaussian function to determine sarcomere length.

For experiments at I22, the sarcomeric X-ray reflections were recorded with a sample-to-detector distance of 8.26 m. After subtraction of the camera background, the diffraction patterns were integrated 0.00304 nm^-1^ on either side of the meridional axis, and the residual background was subtracted as described above. The axial region 5 – 7 µm containing the fourteenth-order reflection was fitted with a single Gaussian peak. Only even orders were observed in this region in resting and contracting muscle, as reported previously (Bordas *et al*., 1987; Hill *et al*., 2021). At ID02 sarcomere reflections up to the twenty-fourth order were visible, confirming the assignment of the 14^th^ order used for sarcomere length determination at I22 (Fig. S1).

### Analysis of Small-Angle X-Ray Diffraction Patterns

The instrumental background was subtracted from the small-angle X-ray diffraction data for each time frame averaged from the series of contractions in each muscle, and each resulting image was centred and aligned using the equatorial 10 reflections via a custom-written automated tilt correction procedure in ImageJ.

#### Equatorial Reflections

The equatorial intensity distribution was determined by integrating from 0.0036 nm^-1^ on either side of the equatorial axis (perpendicular to the muscle axis), and the intensities and spacings of the 10, 11 and 20 reflections were determined by fitting three Gaussian peaks in the radial region 0.02 to 0.065 nm^-1^ with the following constraints:

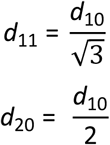

The additional equatorial reflection at ∼0.04 nm^-1^ associated with Z-disc in fast muscle is very weak in rat soleus, so it was not included in the fitting procedure.

The volume of the filament lattice in each sarcomere (V) was calculated as:

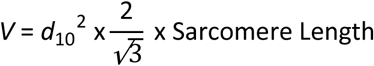

#### Meridional Reflections

To analyse the meridional reflections, aligned 2D patterns were first mirrored horizontally and vertically. The distribution of diffracted intensities along the meridional axis of the diffraction pattern (parallel to the muscle axis) was calculated by integrating from 0.0038 nm^-1^ on either side of the meridian. These integration limits are narrower that those used as in our previous studies of intact mouse EDL (Hill *et al*., 2021, 2022, 2025), and were chosen to provide optimal resolution of the sub-peaks of the M3 reflection with acceptable signal-to-noise ratio (Fig. S2). Intensity distributions were background subtracted using arc hull fitting with smoothing set to 2 using the Baselines extension in Igor Pro. Integrated intensities were obtained for the following axial regions: M3, 0.066–0.072 nm^−1^; M6, 0.135–0.142 nm^−1^. The cross-meridional width of the M3 and M6 reflection was determined by fitting the radial intensity distribution in the axial regions defined above with a double Gaussian function centred on the meridian, taking the narrower fitted width as that of the meridional reflection of interest. The interference components of the M3 and M6 reflections were characterised by fitting multiple Gaussian peaks with the same width to the axial intensity distribution. The additional reflection called the ‘star’ peak that is observed on the low-angle side of the M3 reflection in fast muscle (Caremani *et al*., 2021; Hill *et al*., 2021, 2022) was not resolved in rat soleus muscle. Four additional peaks observed in the reciprocal space region 0.062-0.067 nm^-1^ were fitted with Gaussian functions of equal width; these were not considered to be components of the M3 reflection. The total intensities of the M3 and M6 reflections were calculated as the sum of the intensities of the component peaks of each reflection and multiplied by the cross-meridional width to correct for changes in lateral misalignment between filaments (Huxley *et al*., 1982); the spacing was calculated as the weighted average of that of the component peaks.

#### Spatial Calibration

At I22, reflection spacings were calibrated using an etched silicon-rich nitride grating with a periodicity of 100 nm (Silson, Southam, UK) inserted at the position normally occupied by the muscle. The resulting small-angle X-ray diffraction pattern was aligned and centred as described above and integrated from 0.00083 nm^-1^ either side of the axis containing diffraction peaks corresponding to the 100 nm periodicity. Each peak was fitted with a Gaussian, and reciprocal spacings calibrated in detector pixel units by linear regression of the centres of peaks up to the 11^th^ order on either side of the beamstop.

At ID02, reflection spacings were calibrated using the wavelength of the monochromatic X-ray beam, determined from the atomic absorption edges of several pure elements and accurate to better than 0.01%, the pixel dimensions of the detector, and the position encoder of the detector wagon (Narayanan *et al*., 2018). The limiting factor for calibration of muscle experiments is the ca 1 mm uncertainty in the distance between the muscle and the fixed main flange of the detector tube, but this can be eliminated by measuring the diffraction pattern from the muscle over a range of wagon positions (Brunello *et al*., 2020).

The spacing of the M3 reflection (*S*_M3_) in resting rat soleus muscle determined by these methods was 14.442 ± 0.004 nm (mean ± SD, n = 4) at I22 and 14.452 ± 0.013 nm at ID02 (mean ± SD, n = 7). *S*_M6_ was measured as 7.230 ± 0.002 nm at I22 and 7.226 ± 0.002 nm at ID02. Corresponding values for resting mouse EDL muscle at I22 were recalculated from the data of Hill *et al*. (2021) using the above method as 14.456 ± 0.008 nm for *S*_M3_ and 7.231 ± 0.004 nm for *S*_M6_ (mean ± SD, n = 9), and the time courses of the changes in those spacings in twitch and tetanic contractions are plotted in Fig. 7E and Fig. 8E. These values of *S*_M3_ for resting rat soleus and mouse EDL muscle are not significantly different from that, 14.461 ± 0.013 nm, determined at ID02 using the above calibration method for demembranated rabbit psoas muscle fibres in relaxing solution containing 5% Dextran, 27°C, SL = 2.47 ± 0.05 µm, or that, 14.479 ± 0.007 nm, for intact quiescent trabeculae from rat ventricle, SL = 1.95 µm, 27°C (Brunello *et al*., 2020). However, all the above values for *S*_M3_ are about 1% larger than the 14.34 ± 0.01 nm reported by Haselgrove (1975) for frog sartorius muscle at rest length, which has been used in many subsequent X-ray and electron microscopy studies to calibrate muscle filament periodicities.

#### Off-Axis Layer Line Reflections

The axial intensity distributions of the first myosin and first actin layer lines (ML1 and AL1) were calculated by integrating the radial region between 0.037 nm^-1^ and 0.064 nm^-1^ from the meridional axis, using unmirrored data at I22 to avoid the tile boundaries. For data collected at ESRF, right-left mirroring was used to increase signal:noise. The estimate of the intensity of the ML1 reflection (*I*_ML1_) in Fig. 4E was obtained by integration in the axial region between 0.017 nm^-1^ and 0.024 nm^-1^ to exclude the contribution from the partially overlapped AL1 reflection (Fig. 4A&B, grey shading; Piazzesi *et al*., 1999; Caremani *et al*., 2021). The estimates of the intensities of the off-axial reflections in Figs. 5 and 6 were obtained by integration in the axial regions 0.016–0.034 nm^-1^, 0.041–0.049 nm^-1^ and 0.066–0.072 nm^-1^ to obtain the radial intensity distributions of the first, second and third myosin-based layer lines respectively. Partially mirrored data were used to avoid detector tile boundaries. These radial profiles were then integrated axially in the radial range 0.020-0.035 nm^-1^ corresponding to the equatorial 10 reflection, and 0.037-0.061 nm^-1^ corresponding to the 11 and 20 equatorial reflections (Fig. 5, grey shading).

The optimal radial integration limits for measuring the intensity of the AL1 layer line (*I*_AL1_) were determined by averaging three resting frames and three frames at peak force in the tetanus for each muscle, then averaging data from all the muscles. The layer lines were then radially integrated using a series of 0.01 nm^-1^ wide strips from 0.05-0.06 nm^-1^ to 0.12-0.13 nm^-1^ and background subtracted using arc hull fitting with smoothing set to 2 using the Baselines extension in Igor Pro. The axial spacing of the mixed first layer line (*S*_L1_) was then calculated from the centroid of the distribution in the region 0.0185-0.0355 nm^-1^ (Fig. S3). *S*_L1_ was ∼37.5 nm in the radial region between 0.09 nm^-1^ and 0.12 nm^-1^, as expected for AL1 (Bordas *et al*., 1999).

The intensity of the sixth-order actin layer line (*I*_AL6_) in the ESRF experiments was obtained by integrating fully mirrored data in the radial region 0.035-0.060 nm^-1^ to minimise the contribution of the tails of the meridional-based M7 and M8 reflections, then using global Gaussian deconvolution of the axial region 0.166-0.178 nm^-1^ (Wakabayashi *et al*., 1994; Kiss *et al*., 2018) assuming constant axial spacing and width of the AL6, M7 and M8 reflections. In the experiments at I22 the radial region 0.035-0.057 nm^-1^ was used with unmirrored data to exclude a tile boundary. Gaussian deconvolution could not be used because part of the reflection was off the edge of the detector, so *I*_AL6_ was estimated by integration of the axial region 0.166-0.170 nm^-1^. 1:2:1 smoothing of the time-resolved data was applied for *I*_AL6_ to increase signal-to-noise.

### Correction of Integrated Intensities for changes in muscle mass in the X-ray beam

Small movements of the muscles with respect to the X-ray beam during contraction can alter the mass in the beam and, consequently, the diffracted X-ray intensities (Hill *et al*., 2025). The time courses of X-ray intensity time courses during a tetanus were corrected for this effect by dividing by the relative change in the background intensity under the 1D axial intensity distribution of the ML1/AL1 layer line (between 0.037 nm^-1^ and 0.064 nm^-1^ from the meridian) for each frame with respect to that at rest (Wang *et al*., 2024; Hill *et al*., 2025). The average correction was 10% at the peak of the tetanus. No correction was applied for the twitch, because it would have been less than 1%.

### Statistical Analyses

All statistical analyses were performed using Jamovi (The Jamovi Project, v2.5.6) and Microsoft Excel. Differences in force or X-ray data between the key time periods in the protocol shown in Fig. 1 were analysed using a repeated measures analysis of variance (ANOVA) with Tukey’s post hoc analysis for data where a main effect was observed (Table 1, Table S2, S3). To determine whether non-parametric analyses were required, data were first checked for normality of distribution using Shapiro-Wilks test, skewness, kurtosis and sphericity using Mauchley’s W. Those which were not spherical used a Greenhouse-Geisser sphericity correction. In the event of non-normally distributed data, the non-parametric Friedman’s test with Durbin-Conover pairwise comparison was used. Main effects and post-hoc analyses P-values are provided in Table S2 for tetanus data and Table S3 for twitch data.

**Table 1.** Change in force, sarcomere length and X-ray parameters in the periods of tetanus and twitch defined in Fig. 1A. Three or four time frames were averaged for each period. Mean ± S.E. for n = 4 (tetanus) and n = 7 (twitch). Superscripted letters denote significant differences between key phases using a repeated-measures ANOVA with Tukey’s post hoc analyses. ^a^P<0.05 when compared to rest. ^b^P<0.05 when compared to peak force. ^c^P<0.05 when compared to isometric relaxation (tetanus only).

| Parameter | Tetanus |  |  |  | Twitch |  |  |
| --- | --- | --- | --- | --- | --- | --- | --- |
|  | Rest | Peak Force | Isometric Relaxation | Mechanically Relaxed | Rest | Peak Force | Mechanically Relaxed |
| **Force (kPa)** | 0.65 ± 0.26 | 105.83 ± 23.86<sup>a</sup> | 109.13 ± 23.92<sup>a</sup> | 0.56 ± 0.08<sup>b,c</sup> | 0.65 ± 0.41 | 18.76 ± 2.99<sup>a</sup> | 0.60 ± 0.35 |
| **SL (μm)** | 2.43 ± 0.03 | 2.15 ± 0.04 | 2.14 ± 0.04 | N/A | 2.46 ± 0.05 | 2.42 ± 0.06 | 2.44 ± 0.05 |
| **d<sub>10</sub> (nm)** | 39.10 ± 0.53 | 39.70 ± 0.46<sup>a</sup> | 39.64 ± 0.44<sup>b</sup> | 39.92 ± 0.28<sup>a,b</sup> | 37.86 ± 0.66 | 37.66 ± 0.66 | 38.02 ± 0.63<sup>a</sup> |
| **I<sub>ML1</sub>** | 1 | 0.55 ± 0.25<sup>a</sup> | 0.51 ± 0.23<sup>a</sup> | 0.79 ± 0.18<sup>a,b,c</sup> | 1 | 0.92 ± 0.11<sup>a</sup> | 0.97 ± 0.06 |
| **A<sub>ML1</sub>** | 1 | 0.73 ± 0.16<sup>a</sup> | 0.70 ± 0.16<sup>a</sup> | 0.88 ± 0.11<sup>a,b,c</sup> | 1 | 0.96 ± 0.06<sup>a</sup> | 0.98 ± 0.03 |
| **I<sub>AL1</sub>** | 1 | 1.37 ± 0.50 | 1.45 ± 0.46 | 1.08 ± 0.25 | 1 | 1.41 ± 0.28<sup>a</sup> | 0.91 ± 0.08<sup>a,b</sup> |
| **I<sub>AL6</sub>** | 1 | 1.59 ± 0.40<sup>a</sup> | 1.53 ± 0.28<sup>a</sup> | 0.92 ± 0.17<sup>b,c</sup> | 1 | 1.28 ± 0.32<sup>a</sup> | 1.05 ± 0.12<sup>b</sup> |
| **I<sub>101</sub>** | 1 | 0.60 ± 0.24<sup>a</sup> | 0.49 ± 0.06<sup>a</sup> | 0.83 ± 0.13<sup>c</sup> | 1 | 0.65 ± 0.18<sup>a</sup> | 0.89 ± 0.13<sup>a,b</sup> |
| **I<sub>111+201</sub>** | 1 | 0.39 ± 0.21<sup>a</sup> | 0.40 ± 0.23<sup>a</sup> | 0.84 ± 0.15<sup>a,b,c</sup> | 1 | 0.83 ± 0.06<sup>a</sup> | 0.96 ± 0.03<sup>a,b</sup> |
| **I<sub>M6</sub>** | 1 | 0.77 ± 0.29<sup>a</sup> | 0.74 ± 0.23<sup>a</sup> | 0.83 ± 0.23<sup>c</sup> | 1 | 0.80 ± 0.16<sup>a</sup> | 0.89 ± 0.04<sup>a,b</sup> |
| **S<sub>M6</sub> (nm)** | 7.230 ± 0.002 | 7.292 ± 0.015<sup>a</sup> | 7.294 ± 0.014<sup>a</sup> | 7.232 ± 0.002<sup>b,c</sup> | 7.226 ± 0.002 | 7.242 ± 0.007<sup>a</sup> | 7.228 ± 0.001<sup>a,b</sup> |
| **I<sub>M3</sub>** | 1 | 0.87 ± 0.27 | 0.86 ± 0.25 | 0.76 ± 0.14<sup>a</sup> | 1 | 0.52 ± 0.10<sup>a</sup> | 0.89 ± 0.03<sup>a,b</sup> |
| **A<sub>M3</sub>** | 1 | 0.93 ± 0.14 | 0.92 ± 0.13 | 0.87 ± 0.08<sup>a</sup> | 1 | 0.72 ± 0.07<sup>a</sup> | 0.94 ± 0.02<sup>a,b</sup> |
| **S<sub>M3</sub> (nm)** | 14.442 ± 0.004 | 14.558 ± 0.039<sup>a</sup> | 14.561 ± 0.039<sup>a</sup> | 14.438 ± 0.004<sup>b,c</sup> | 14.452 ± 0.013 | 14.446 ± 0.015<sup>a</sup> | 14.450 ± 0.012<sup>a</sup> |
| **I<sub>11</sub>/I<sub>10</sub>** | 0.58 ± 0.04 | 1.36 ± 0.33<sup>a</sup> | 1.37 ± 0.33<sup>a</sup> | 0.56 ± 0.06<sup>b,c</sup> | 0.48 ± 0.03 | 0.62 ± 0.05<sup>a</sup> | 0.49 ± 0.03<sup>a,b</sup> |

Paired-sample T-tests were used to determine whether the half-time and rate constants for X-ray data differed significantly from those of force (Table 2, Table S4). When data were not normally distributed, as determined by checks for normality of distribution by Shapiro-Wilks test, skewness, and kurtosis, the non-parametric Wilcoxon’s signed-rank test was used.

**Table 2.** Half-times and rate constants of force, sarcomere length and X-ray signals. Activation and relaxation half-times (*t*_1/2_) were derived from single sigmoidal fits except for *I*_M3_, *A*_M3_ and *S*_M3_ tetanus activation, where a double sigmoidal function was used to obtain *t*_1/2_ for the descending (Fast) and ascending (Slow) phases. Tetanus and twitch activation data were fit between -26 ms and 234 ms and -17.75 ms and 52.25 ms, respectively, and *t*_1/2_ is given with respect to *t* = 0 ms (i.e. the first stimulus). Tetanus relaxation data were fit between 234 ms and 604 ms and *t*_1/2_ given with respect to the last stimulus. Twitch relaxation data were fit between 52.25 ms and 327.75 ms and *t*_1/2_ given with respect to *t =* 52.25 ms. Rate constants for isometric relaxation (slow *K*_REL_) in the tetanus was calculated as the slope of the linear fit between 244 ms and 324 ms normalised by the difference between rest and the first frame after the last stimulus (244 ms). Rate constants for the subsequent exponential relaxation (fast *K*_REL_) were calculated by exponential fits from 344 ms to 604 ms. N/A denotes parameters that could not be obtained by the above fitting approaches. Values presented as mean ± S.D. for n = 4 muscles for tetanus and n = 7 for twitch; ^a^n = 3. *P<0.05 when comparing a structural parameter against force for half-times and rate constants using Student’s paired-samples *t-*test.

| Parameter | Tetanus |  |  |  | Twitch |  |
| --- | --- | --- | --- | --- | --- | --- |
| | Tetanus Activation<br>$t_{1/2}$ (ms) | Tetanus Relaxation<br>$t_{1/2}$ (ms) | Slow $K_{REL}$ (s<sup>-1</sup>) | Fast $K_{REL}$ (s<sup>-1</sup>) | Twitch Activation<br>$t_{1/2}$ (ms) | Twitch Relaxation<br>$t_{1/2}$ (ms) |
| Force | 72.3 ± 9.8 | 121.6 ± 4.7 | 1.7 ± 0.3 | 23.0 ± 1.9 | 18.0 ± 1.6 | 28.2 ± 3.9 |
| $I_{ML1}$ | 72.7 ± 38.0 | 123.5 ± 26.3<sup>a</sup> | 1.7 ± 0.5 | 11.1 ± 8.6 | N/A | N/A |
| $A_{ML1}$ | 81.3 ± 32.1 | 124.7 ± 26.7<sup>a</sup> | 2.2 ± 0.7 | 19.6 ± 11.7 | N/A | N/A |
| $I_{111+201}$ | 47.9 ± 6.1* | 207.7 ± 43.5* | 0.9 ± 0.4 | 8.8 ± 1.1* | 24.1 ± 6.4 | 55.5 ± 5.7* |
| $S_{M6}$ | 44.2 ± 16.7* | 140.2 ± 14.6 | 2.5 ± 1.3 | 12.3 ± 1.2* | 16.5 ± 8.9 | 31.2 ± 16.3 |
| $I_{M3}$ | Fast – 23.0 ± 12.9*<br>Slow – 112.8 ± 43.2 | N/A | N/A | N/A | 9.9 ± 2.6* | 89.4 ± 18.1* |
| $A_{M3}$ | Fast – 28.9 ± 12.1*<br>Slow – 106.0 ± 24.9 | N/A | N/A | N/A | 13.6 ± 2.4* | 91.9 ± 21.2* |
| $S_{M3}$ | Fast – 17.1 ± 9.0<sup>a</sup><br>Slow – 78.3 ± 10.2 | 101.7 ± 12.4 | 4.3 ± 2.2 | 53.4 ± 24.5* | N/A | N/A |
| $I_{11}/I_{10}$ | 61.0 ± 22.6 | 114.3 ± 9.7 | 4.3 ± 0.8* | 17.4 ± 2.7* | 15.7 ± 3.3* | 41.7 ± 10.0* |

Data presented are mean ± SD throughout. Significance was set at P<0.05 for all analyses.

## Results

Rat soleus muscles were electrically stimulated at 27°C to produce a twitch or a short, fused tetanus at constant muscle length (Fig. 1A). Peak force in the tetanus was about 110 kPa (Table 1), about half of that typically produced at the plateau of a tetanus in fast mammalian muscle at the same temperature (Percario *et al*., 2018; Caremani *et al*., 2019; Hill *et al*., 2021). Force continued to rise for about 30 ms after the last stimulus, then declined at an increasing rate for about 100 ms (Fig. 1A, yellow shading), when it became roughly exponential, with rate constant 23 s^-1^ (Table 2). We refer to the pre-exponential phase of relaxation as ‘isometric relaxation’ (blue shading), because sarcomere length is constant in this period, as described below. Peak force in the twitch, which was only about 15% of that in the tetanus, was attained about 50 ms after the stimulus, and the exponential phase of relaxation in the twitch had a half time of 28 ms (Table 2).

We determined changes in the structure of the contractile filaments during the twitch and tetanus using small-angle X-ray diffraction, exploiting the almost crystalline order of the contractile filaments in the muscle sarcomere. The myosin-containing thick filaments (Fig. 1B, larger open circles) and actin-containing thin filaments (smaller filled circles) are arranged in a hexagonal lattice, and the filaments themselves are helical (Fig. 1C). The periodic repeats of these structures give rise to characteristic features in the X-ray diffraction pattern from a muscle mounted with its long axis vertical (Fig. 1D). Two relatively bright spots on the horizontal axis of the pattern, the 10 and 11 equatorial reflections (Fig. 1D, green label), can be thought of as reflections of the incident X-ray beam by the 10 and 11 planes of the hexagonal lattice (Fig. 1B). The 10 planes are separated by a distance *d*_10_, and define the unit cell of the hexagonal lattice, passing through the centres of the thick filaments (Fig. 1B).

The axial periodicities of the filaments (Fig. 1C) produce X-ray reflections on the vertical axis of the diffraction pattern called meridional reflections (Fig. 1D, magenta labels). The dominant reflections are associated with the ca 43 nm helical periodicity of the thick filament (Fig. 1C, *S*_ML1_) and labelled M1, M2, M3 etc as orders of that fundamental repeat. The M3 reflection, corresponding to the ca 14.4 nm axial periodicity of the three crowns of myosin motors in each 43 nm repeat (Fig. 1C, *S*_M3_) is particularly strong (Fig. 1D). Weaker off-axis reflections (Fig. 1D, white labels) are observed with the same separation from the equator as the meridional reflections, and the same separation from the meridian as the equatorial reflections. These off-axial reflections are labelled xyz, where x and y denote the equatorial reflection in the same vertical column, and z denotes the meridional reflection in same horizontal row. The presence of these discrete off-axial reflections shows that the three pairs of myosin motors at each axial level of the thick filament (Fig. 1B, blue ovals) are azimuthally aligned in rat soleus muscle (Ma *et al*., 2019; Gong *et al*., 2022), an arrangement first described in bony fish muscle (Harford & Squire, 1986; Squire *et al*., 2004). This arrangement contrasts with that in most fast muscles of vertebrates, which have azimuthally disordered thick filaments and produce diffraction patterns with a series of smooth ‘layer lines’ parallel to the equator and aligned with the M series of meridional reflections (Huxley & Brown, 1967). The discrete off-axial X-ray reflections produced by rat soleus muscles, sometimes called ‘sampled layer lines’, provide additional information about the conformation of the myosin motors (Koubassova *et al*., 2022).

### Sarcomere length changes

Although the muscle length was held constant in these experiments, the sarcomeres in the central region of the muscles shorten during active force development as the tendons are stretched. We measured the average sarcomere length in the central region of the muscle illuminated by the X-ray beam using the sarcomere-based X-ray reflections recorded at very low diffraction angles (Fig. 1D,E, red inset). This information cannot be obtained by conventional diffraction or imaging with visible light in thick muscles like rat soleus because visible light is scattered many times as it passes through the muscle, whereas X-rays are scattered only once. The X-ray optics used for the experiments with tetanic stimulation at I22 (Diamond Light Source, UK) allowed us to record the 14^th^ to the 24^th^ even-order reflections of the sarcomere repeat (Fig. 1D, inset; Fig. 2A). The changes in the positions of these reflections showed that sarcomeres in the central region of the muscles shortened smoothly from 2.43 ± 0.03 µm at rest to 2.15 ± 0.04 µm at peak force in the tetanus (Fig. 2C, filled symbols; Table 1) with a half-time of 29.1 ± 5.3 ms, much faster than that of force, indicating that the tendons are more compliant at lower force. There was no significant change in sarcomere length during isometric relaxation (Fig. 2C, blue shading). It was not possible to measure sarcomere length during exponential relaxation because the appearance of multiple sarcomere populations with different lengths made it impossible to assign the high order sarcomere-based reflections (Hill *et al*., 2025).

**Figure 2.**
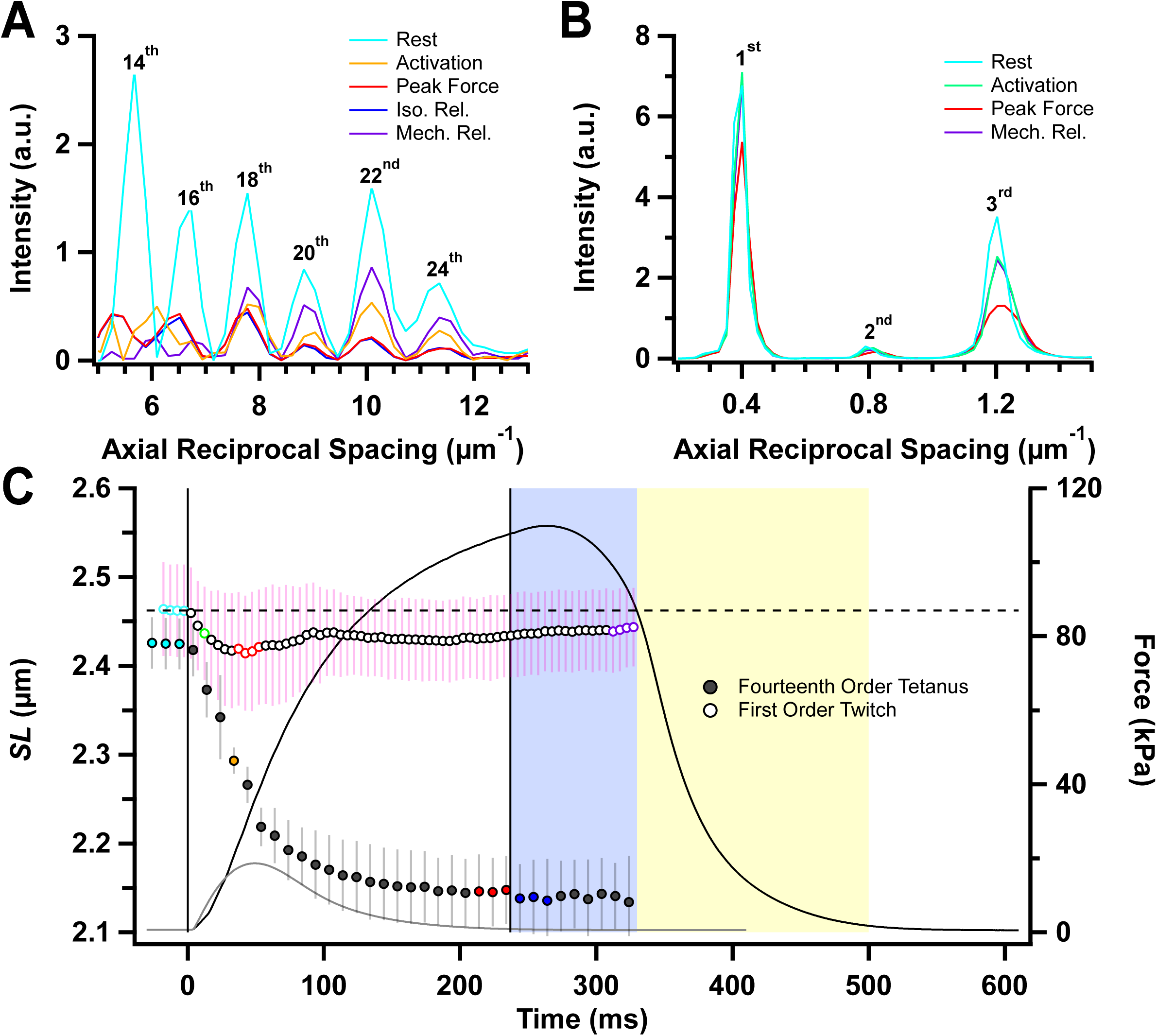
Sarcomere reflections and sarcomere length changes. Ultra-small-angle meridional intensity distribution showing sarcomere reflections collected during the tetanus (*A*) and twitch (*B*) after subtraction of the diffuse background, with orders of sarcomere reflections labelled. *C*, Sarcomere length (*SL*) (mean ± SD) determined from the centroid of the first- and fourteenth-order sarcomere reflections in twitch (open symbols, n = 7 muscles) and tetanus (filled symbols, n = 4 muscles), respectively, with corresponding mean force time course. Vertical solid lines in *C* denote the first and last electrical stimulus. Horizontal dashed line in *C* denote resting *SL* value for the twitch. Colour coding denotes periods shown in Fig. 1A.

The sarcomere length changes during twitches were measured at beamline ID02 at the ESRF, which allowed the distance between the muscle and the X-ray detector to be increased to about 31 m, so that many orders of the sarcomere reflections including the first order could be recorded with high precision (Fig. 1E inset; Fig. S1, black). The resting sarcomere length was slightly longer in the batch of muscles used for the twitch experiments, and decreased by less than 2% following stimulation, with incomplete recovery during mechanical relaxation (Fig. 2C, open circles).

### The lattice of thick and thin filaments

The hexagonal lattice of thick and thin filaments (Fig. 1B) produces a series of equatorial X-ray reflections (Fig. 3A,B) of which the strongest are the 10, associated with the planes of thick filaments bordering the hexagonal unit cell, and the 11, corresponding to the planes passing through the centres of the thick filaments and the thin filaments. An additional X-ray reflection of unknown origin was observed at a lower angle than the 10, corresponding to a periodicity of about 71 nm (Fig. 3A). This inner equatorial reflection has not been reported in diffraction patterns from fast skeletal muscle of either amphibians or mammals. The ‘z’ equatorial reflection from the square lattice of thin filaments at the Z band of the sarcomere, which is present in fast mammalian muscles, is very weak in rat soleus (Ma *et al*., 2019).

**Figure 3.**
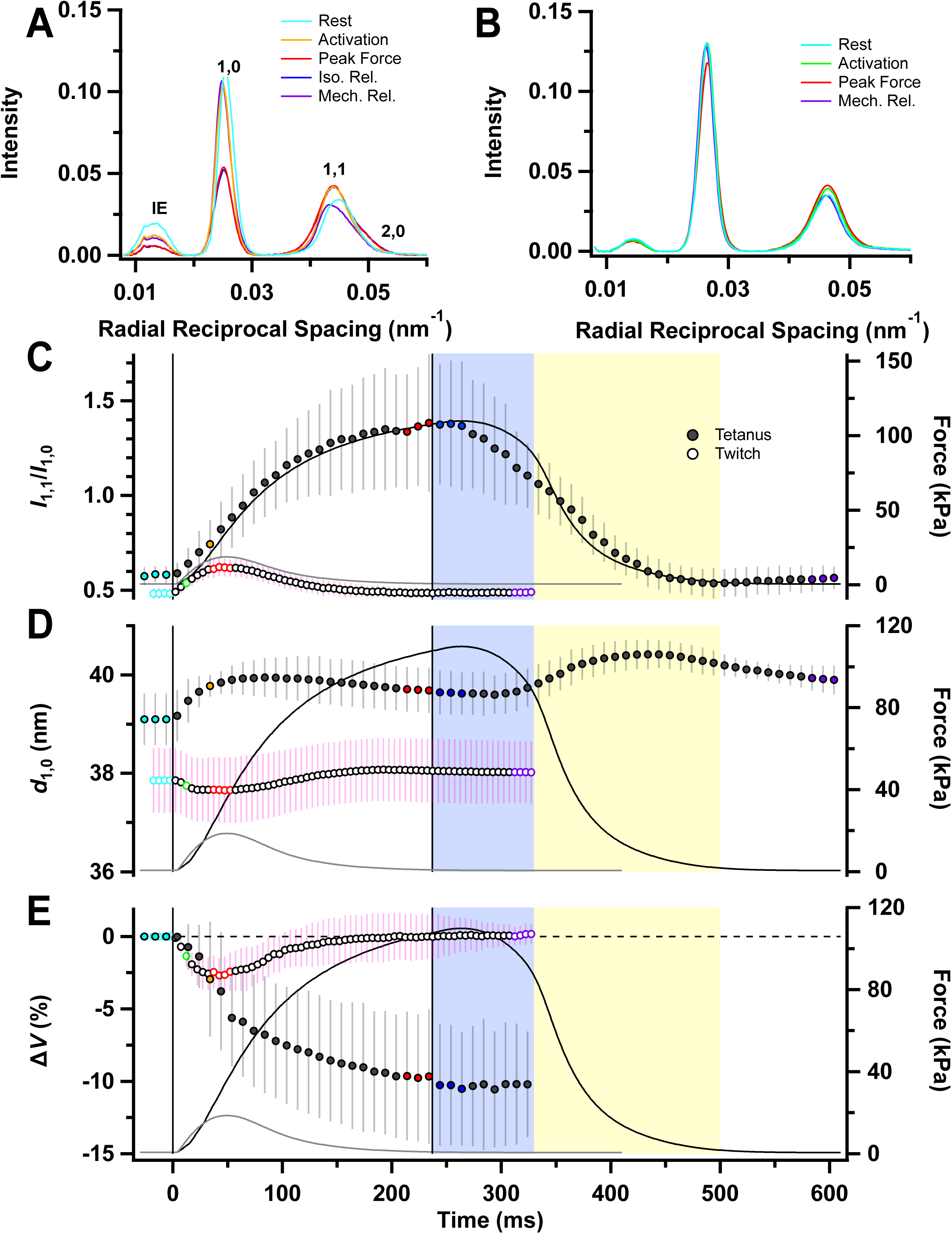
Equatorial reflections. *A* and *B*, Radial intensity distributions along the equator, perpendicular to the long axis of muscle, in tetanus (*A*) and twitch (*B*), with 10, 11, 20 and inner equatorial (IE) reflections labelled in *A*. *C*-*E*, Equatorial intensity ratio (*I*_11_/*I*_10_, *C*), lattice spacing (*d*_10_, *D*) and filament lattice volume (Δ*V*) as a percentage of resting volume (*E*) for tetanus (filled symbols; mean ± SD, n = 4 muscles) and twitch (open symbols; n = 7 muscles). Vertical solid lines in *C-E* denote the first and last electrical stimulus. Horizontal dashed line in *E* denote resting Δ*V* value for the twitch. Colour coding denotes periods shown in Fig. 1A.

The intensity of the 10 reflection (*I*_10_) decreases during active contraction of rat soleus muscles, and that of the 11 reflection (*I*_11_) increases. The intensity ratio (*I*_11_/*I*_10_), which is widely used as an index of the movement of myosin motors from the vicinity of the surface of the thick filaments towards the thin filaments (Haselgrove & Huxley, 1973), was about 0.5 in resting muscles (Fig. 3C; Table 1). Resting *I*_11_/*I*_10_ was slightly lower in the batch of muscles used for the twitch experiments, probably because of the longer sarcomere length. It increased to about 1.4 at peak force in the tetanus (Table 1), with a time course similar to that of force development (Table 2). This peak value was maintained for about 40 ms after the last stimulus, then *I*_11_/*I*_10_ started to decline roughly linearly at about 4 s^-1^, continuing at about the same rate into the first ca 50 ms of exponential relaxation. The increase in *I*_11_/*I*_10_ was much smaller in a twitch, with a peak value of only 0.62 ± 0.04 in comparison with 0.48 ± 0.03 at rest in that batch of muscles, and a time course similar to that of force.

The separation between the 10 planes of the filament lattice (*d*_10_; Fig. 1C) was lower in the resting muscles used in the twitch experiments (Fig. 3D), as expected from their slightly longer sarcomere length. *d*_10_ increased slightly during force development in a tetanus, then again during exponential relaxation, so that the last recorded value after full mechanical relaxation was higher than that at rest. *d*_10_ decreased slightly and transiently in the twitch.

The volume of the sarcomere lattice in the region of the muscles in the X-ray beam decreased by about 12% during force development in the tetanus (Fig. 3E, filled circles; Table 1). The decrease in lattice volume associated with sarcomere shortening is not compensated by lateral expansion, suggesting that myosin motors contribute a lateral compressive force during contraction. The volume decrease was about 3% during the twitch (Fig. 3E, open circles).

### Off-axial reflections

The intensities of the off-axial reflections give information about the mass associated with helical periodicities of the thick and thin filaments (Fig. 1C), and have been used extensively in previous X-ray studies of fast skeletal muscle to follow the transfer of mass of the myosin motors during force development from the helically-ordered thick filament OFF state to take up the distinct helical periodicity of the thin filament in the strongly attached force-generating state. This transition is expected to produce a decrease in the intensity of the ca 43 nm off-axial ‘myosin first layer line’ (*I*_ML1_), and an increase in that of the ca 38 nm ‘actin first layer line’ (*I*_AL1_). These changes are much smaller in rat soleus than in the fast muscles used in previous studies, as indicated by the lack of change in the profile of the mixed layer line on activation (Fig. 4A,B). Moreover, the myosin-based layer lines are strongly sampled in rat soleus muscles, appearing as discrete off-axis reflections (Fig. 1D). The intensities of the individual reflections along a myosin-based layer line may have different time courses during muscle activation, so that the measured intensity change or its time course depends on the radial region used for the measurement.

**Figure 4.**
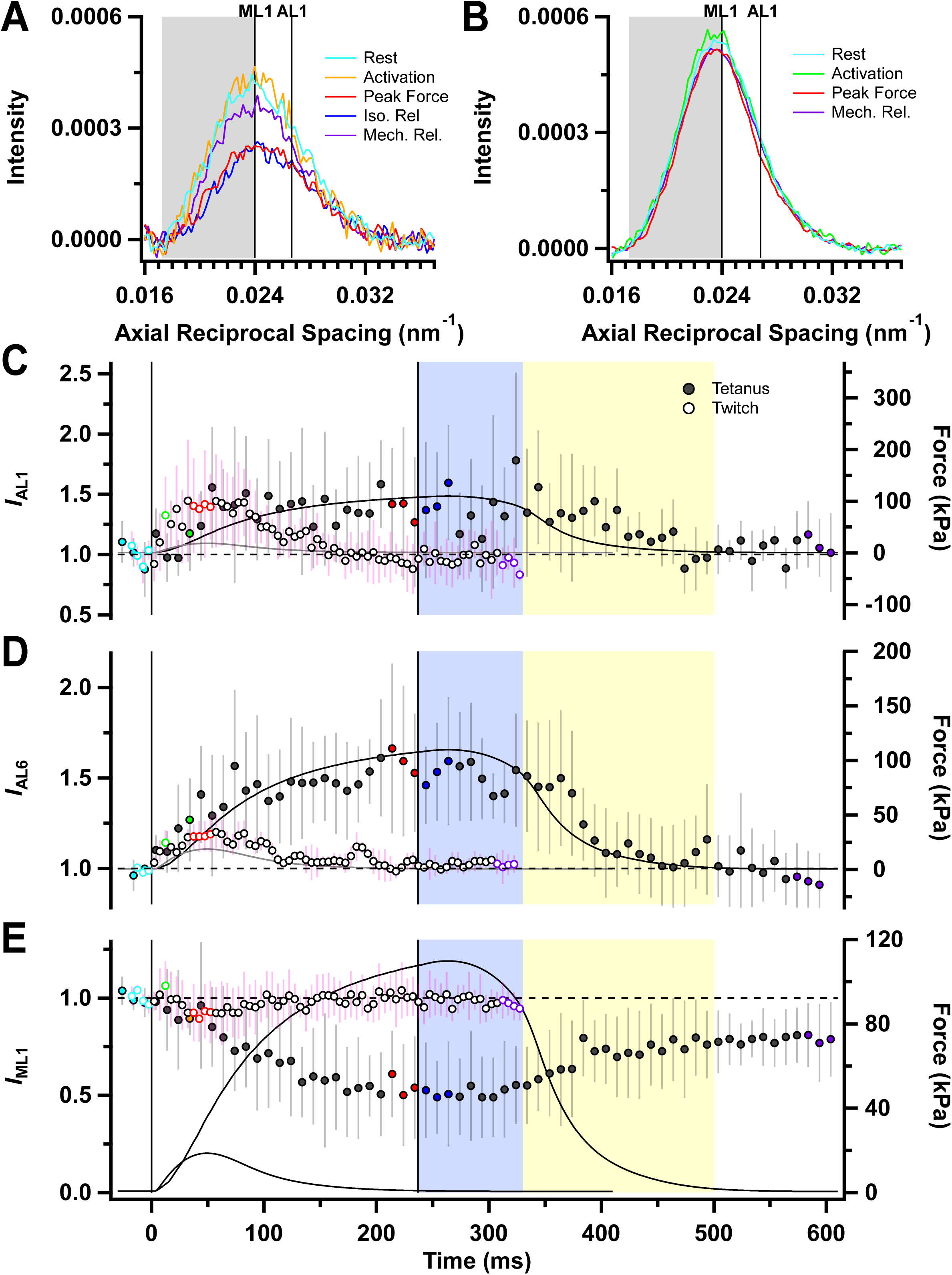
Layer-line reflections. *A* and *B*, Axial intensity distributions of layer line reflections in tetanus (*A*) and twitch (*B*), calculated as described in the text. *C*-*E*, intensities of the first-order myosin layer line (*I*_ML1_, *C*), first-order actin layer line (*I*_AL1_, *D*) and the sixth-order actin layer line (*I*_AL6_, *E*) for tetanus (filled symbols; mean ± SD, n = 4 muscles) and twitch (open symbols; mean ± SD, n = 7 muscles except for *I*_AL6_ which is n = 3), normalised to their respective resting intensities. Grey shading in *A* and *B* corresponds to the lower-angle axial region which were integrated to yield Fig. 4E. Vertical solid lines in *C-E* denote the first and last electrical stimulus. Horizontal dashed line in C-*E* denote resting values. Colour coding denotes periods shown in Fig. 1A.

Given those constraints, we first estimated the change in *I*_AL1_ in the present experiments at high radius, where the myosin-based layer lines are very weak (see Methods for details), as shown by the axial spacing of the mixed layer line in that region, which was 37.3 ± 0.8 nm at rest and 37.6 ± 0.7 nm at peak force in the tetanus, consistent with values expected for a pure AL1 layer line. *I*_AL1_ in this region increased by about 40% at peak force in both the twitch and the tetanus (Fig. 4C; Table 1), although the signal-to-noise ratio is low because AL1 is relatively weak in this region. The intensity of the sixth actin-based layer line (*I*_AL6_), which does not overlap with any myosin-based layer line, also increased by about 40% in the tetanus (Fig. 4D) and by about 20% in the twitch (Table 1).

We used two different methods to estimate the changes in the intensity of the myosin-based off-axial reflections. First, we determined the diffracted intensity in the region conventionally associated with the first myosin layer line in fast muscle by integrating the low-angle half of the mixed axial first order layer line profile in a radial region around the 11 equatorial reflection where AL1 is expected to make little contribution (Fig. 4A,B, grey shading; see Methods), which we will call “*I*_ML1_”. This decreased to about 50% of its resting value during force development in the tetanus (Fig. 4E, filled circles), but retained more than 90% of its resting value in the twitch (open circles). Secondly, we determined the intensities of the individual off-axis reflections from the radial intensity profiles of the first three myosin-based layer lines (Fig. 5), which show a series of peaks at radial positions corresponding to the 10 and 11 equatorial reflections (Fig. 5A,B; vertical dashed lines), with weaker contributions from higher order reflections. The radial profiles of the meridional M1, M2 and M3 reflections, centred on zero radial spacing, are included for comparison. In resting muscle (Fig. 5, cyan), the 111 peak is stronger than the 101, and the 112 is stronger than the 102 on the ML2 layer line, but on the ML3 layer line the 103 is stronger than the 113. The same pattern of relative intensities has been reported for plaice fin muscle (Squire *et al*., 2004; Koubassova *et al*., 2022), and can be reproduced by simple models of helical myosin filaments only if the helix is perturbed, for example if one out of every three layers of myosin motors is disordered, as observed in cryo-EM images of isolated myosin filaments and myosin filaments in myofibrils from cardiac muscle (Tamborrini *et al*., 2023; Dutta *et al*., 2023; Koubassova *et al*., 2025).

**Figure 5.**
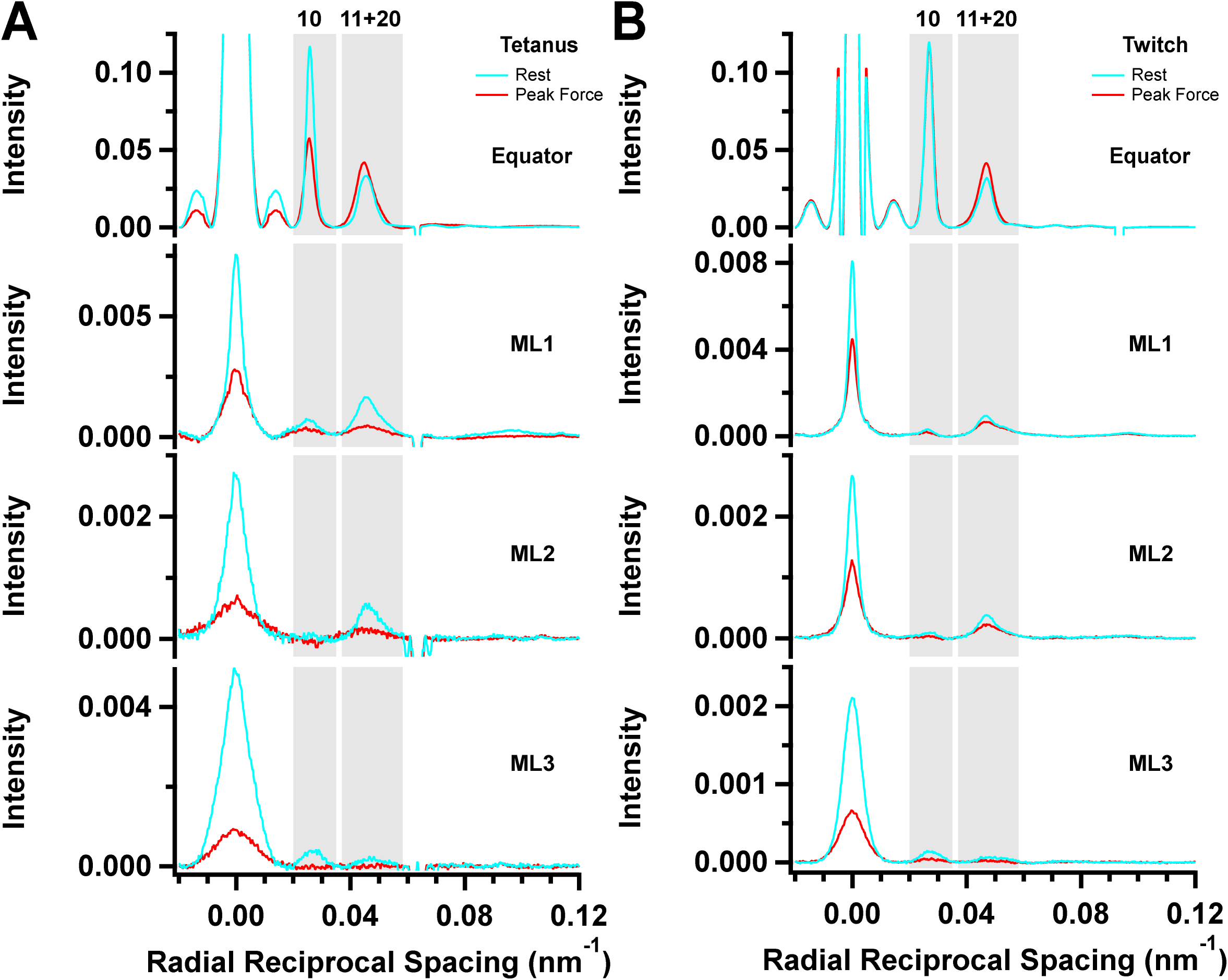
Radial intensity distributions of the myosin-based layer lines. Intensity distributions along the first-, second- and third-order myosin layer lines reflection (ML1, ML2 and ML3, respectively) in tetanus (*A*, averaged from n = 4 muscles) and twitch (B, averaged from n = 7 muscles). Cyan, resting muscle; red, peak force. Corresponding distributions along the equator are shown for comparison. Grey shading corresponds to the regions along the radial intensity distributions which were integrated in the positions of the 10 and 11+20 equatorial-based reflections to yield Fig. 6.

These off-meridional reflections are generally weaker at the peak of the tetanus (Fig. 5A, red) than at rest (cyan), but the fractional intensity change is different for the different reflections. Two peaks could be resolved on the first myosin layer line, corresponding to the 101 reflection and an unresolved mixture of the 111 and 201 reflections. The intensity of the 101 reflection (*I*_101_) decreased to about 70% of its resting value at the peak of either a tetanus or twitch (Fig. 6A; Table 1), with a time course faster than that of force development in the tetanus (Table 2). The combined intensity of the 111 and 201 reflections (*I*_111+201_, Fig. 6B; Table 1) decreased to about 40% of the resting value at peak force in the tetanus, with a time course similar to that of force, and decreased to only about 80% in a twitch. There was no significant recovery of either *I*_101_ or *I*_111+201_ during isometric relaxation from the tetanus (blue shading), and recovery was still incomplete at mechanical relaxation.

**Figure 6.**
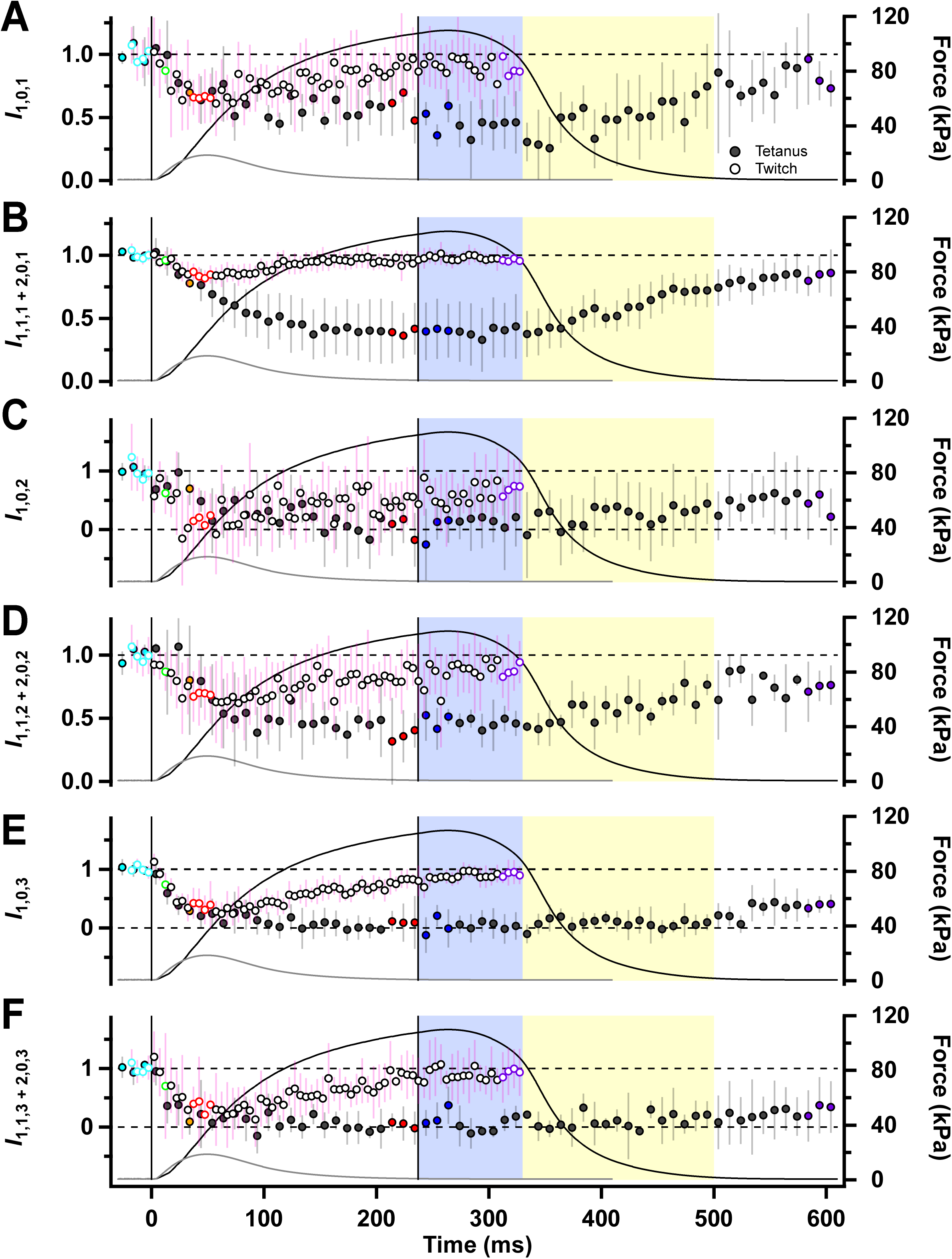
Myosin-based off-axial reflections. Intensities of the 101 (*I*_101_; *A*), mixed 111 and 201 (*I*_111+201_; *B*), 102 (*I*_102_; *C*), mixed 112 and 202 reflections (*I*_112+202_; *D*), 103 (*I*_103_; *E*) and mixed 113 and 203 reflections (*I*_113+203;_ *F*) for tetanus (filled symbols, mean ± SD, n = 4 muscles) and twitch (open symbols, n = 7 muscles), normalised to the respective resting values. Vertical solid lines in *A-F* denote the first and last electrical stimulus. Horizontal dashed line in A-*F* denote either resting values at 1 or minima values at 0. Colour coding denotes periods shown in Fig. 1A.

*I*_102_ is weak in resting muscle (Fig. 5A) and was not detectable at the peak of either a twitch or tetanus (Fig. 6C). *I*_112+202_ (Fig. 6D) is stronger at rest and decreases during a twitch or tetanus in roughly the same way as *I*_111+201_. *I*_103_ (Fig. 6E) and (*I*_113+203_; Fig. 6F) also became very weak during either a twitch or tetanus, with a time course faster than that of force development in the tetanus. Quantitative interpretation of these intensity changes in relation to that of the conventional “*I*_ML1_” will require a structural model of the arrangement of the myosin motors on the thick filament. Qualitatively, they show a striking loss of the helical arrangement of the myosin heads characteristic of relaxed muscle that is much faster than force development in the tetanus, and which is observed at the much smaller peak force in the twitch. We return to the interpretation of these unexpected findings in the Discussion.

### Meridional reflections

The M3 meridional reflection is associated with the axial periodicity of crowns of myosin motors along the thick filaments. In resting muscle, it appears as a strong peak corresponding to an axial periodicity of about 14.44 nm (Fig. 7A,B, cyan, MA peak), with a shoulder on the high-angle side (HA peak) with axial periodicity about 14.23 nm. A third peak (Fig. 7A, red, LA peak) becomes prominent in a tetanus, with axial periodicity about 14.69 nm. The relative intensities of these three peaks are similar to those reported previously in fast muscles and muscle fibres (Linari *et al*., 2000; Hill *et al*., 2021; Caremani *et al*., 2023). Their spacings are roughly 1% larger than those reported previously for fast muscle, but this difference is due to our use of an absolute calibration of the small-angle X-ray beamlines, in contrast with the previous assumption that the spacing of the M3 reflection is 14.34 nm (Haselgrove, 1975; see Methods). A smaller additional peak, M3*L*, corresponding to an axial periodicity of about 15.52 nm, is present in resting rat soleus muscles, and may be the third order of the long or *L* periodicity also seen in intact (Hill *et al*., 2021) and demembranated (Caremani *et al*., 2021) fast mammalian muscle fibres. The additional ‘star’ peak seen in fast muscle (Caremani *et al*., 2021; Hill *et al*., 2021), corresponding to an axial periodicity of about 15.0 nm, was not detected in rat soleus muscles.

The total integrated intensity of the M3 reflection (*I*_M3_), determined as the sum of the intensities of the LA, MA and HA peaks, decreases transiently at the start of a tetanus (Fig. 7C, filled circles), reaching a minimum after about 50 ms. The early decrease in *I*_M3_ is likely to be associated with the loss of the folded helical motors characteristic of resting muscle, and the later increase with the slower formation of actin-attached force-generating motors with their long axes roughly perpendicular to the filament axis (Irving *et al*., 2000; Hill *et al*., 2021). At the end of electrical stimulation, *I*_M3_ starts to decrease during isometric relaxation (blue shading), suggesting that motors are already detaching from actin during that period, as also suggested by the accompanying decrease in the equatorial intensity ratio (*I*_11_/*I*_10_; Fig. 3C). *I*_M3_ does not fully recover its resting value at mechanical relaxation, suggesting that the folded helical OFF conformation of the myosin motors recovers more slowly than force relaxation, as also signalled by *I*_ML1_ (Fig. 4E) and *I*_111+201_ (Fig. 6B). Surprisingly, the time course of *I*_M3_ in a twitch (Fig. 7C, open circles) is indistinguishable from that in a tetanus (filled circles), although the force is much smaller and the fine structure of the M3 reflection does not change in the twitch (Fig. 7B).

The spacing of the M3 reflection (*S*_M3_; Fig. 7D) decreases slightly for the first 20 ms of the tetanus suggesting that, like *I*_M3_, it contains a fast, decreasing phase followed by a slow increase (Table 2). *S*_M3_ also decreases during isometric relaxation (blue shaded area) and is lower than its resting value during late exponential relaxation. *S*_M3_ decreases slightly during a twitch (Fig. 7D, open circles), with no sign of the increasing component seen in a tetanus.

These changes in *S*_M3_ are strikingly different from those observed in fast muscle, in both kinetics and amplitude (Fig. 7E; Hill *et al*., 2021). The mean half-time of the increase in *S*_M3_ during force development in the tetanus in rat soleus, 78 ms (Table 2), is about seven times slower than that in mouse EDL muscle at the same temperature (11 ms; Hill *et al*., 2021), compared with a roughly four-fold difference in the half-time for force development. The amplitude of the increase in *S*_M3_ in a tetanus in rat soleus is about half that seen in mouse EDL. *S*_M3_ decreases in a soleus twitch, but *increases*, to about half the tetanus plateau value, in EDL.

**Figure 7.**
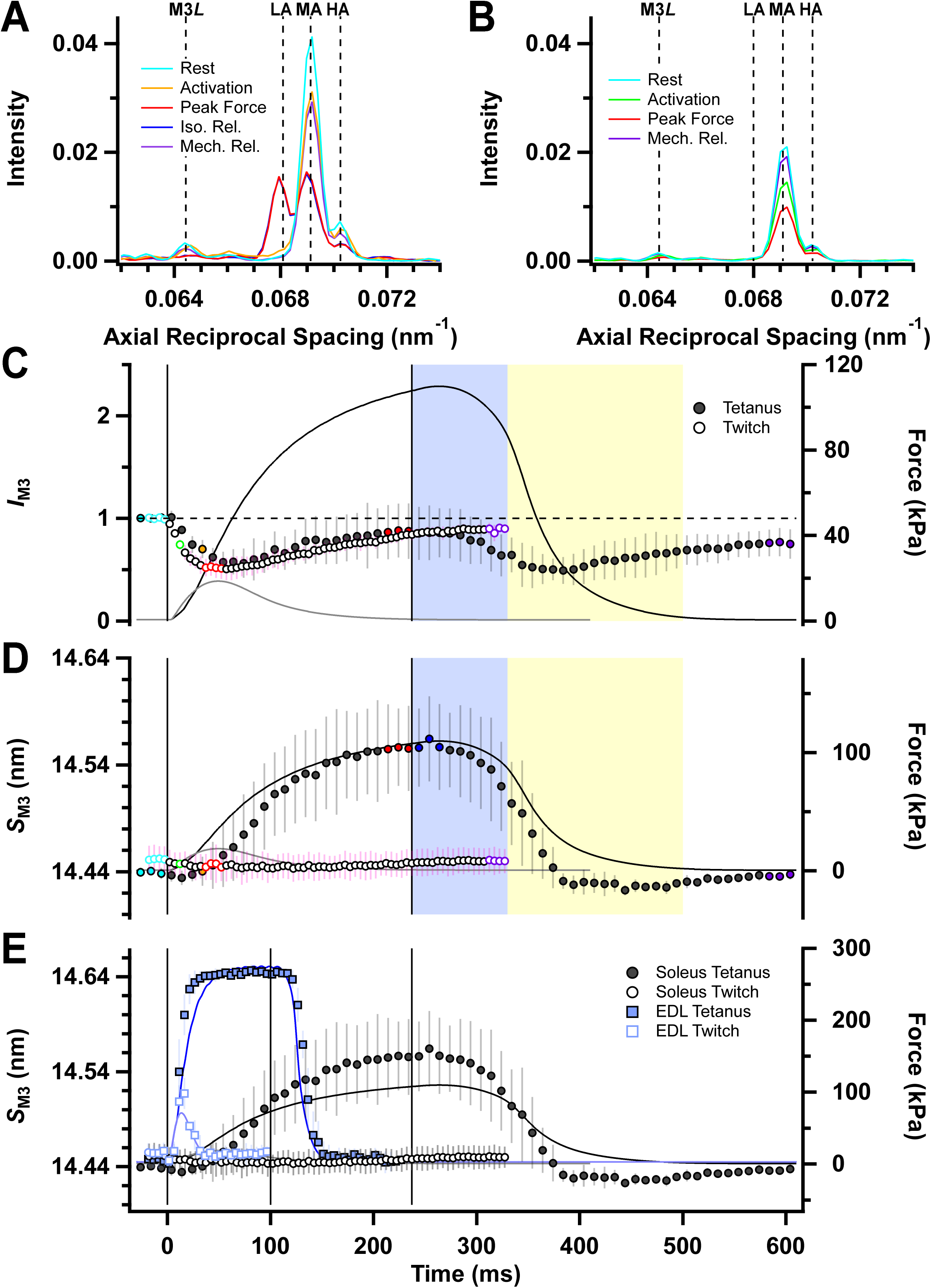
Myosin-based M3 reflection. *A* and *B*, Axial intensity distributions of the M3 reflection in tetanus (*A*) and twitch (*B*). Vertical dashed lines denote low-angle (LA), mid-angle (MA), high-angle (HA) peaks of the M3, and M3*L* reflection. Colour coding denotes periods shown in Fig. 1A. *C* and *D*, Intensity of the M3 reflection (*I*_M3_, *C*) normalised to its resting value, and spacing of the M3 reflection (*S*_M3_, *D*) for tetanus (filled symbols, mean ± SD, n = 4 muscles) and twitch (open symbols, n = 7 muscles) protocols. In *E*, *S*_M3_ for the soleus (circles) is compared against *S*_M3_ from fast-twitch mouse EDL muscle (blue) during the twitch (open squares, mean ± SD, n = 4 muscles) or 100 ms tetanus (filled squares, n = 5 muscles). *S*_M3_ and force (blue traces) in mouse EDL is from Hill *et al*. (2021), but have been re-calculated using the beamline calibration for those data. Intensities in *A*, *B* and *C* corrected for changes in radial width. Vertical solid lines in *C-E* denote the first and last electrical stimulus for each muscle type. Horizontal dashed line in *C* denote resting *I*_M3_. Colour coding denotes periods shown in Fig. 1A.

The M6 meridional reflection has a spacing of about 7.23 nm in resting soleus muscle. Three sub-peaks can be resolved, designated LA, MA and HA (Fig. 8A,B). An additional weaker peak on the low angle side of the main reflection, with an axial periodicity of about 7.62 nm, corresponds to the M6*L* reflection (Caremani *et al*., 2021; Hill *et al*., 2021). The total intensity of the M6 reflection (*I*_M6_) decreased during both twitch and tetanus (Fig. 8C, red), in contrast with the lack of change in fast mammalian muscle (Hill *et al*., 2021). Its spacing (*S*_M6_; Fig. 8D) increased monotonically in both twitch and tetanus, with no evidence of the early decrease seen in *S*_M3_. *S*_M6_ reproducibly *increases* in a twitch, in marked contrast to the decrease in *S*_M3_*. S*_M6_ increased faster than force in the tetanus (Table 2), and this increase was three times faster than that of *S*_M3_. These changes in *S*_M6_ are distinct from those seen in fast muscle, in both kinetics and amplitude (Fig. 8E). The mean half-time of the increase in *S*_M6_ in a tetanus in rat soleus, 44 ms (Table 2), is about five times slower than that in mouse EDL muscle (8 ms; Hill *et al*., 2021), corresponding to the roughly four-fold difference in the half-time for force development. The increase in *S*_M6_ during a tetanus in rat soleus is about half that seen in mouse EDL, similar to the relative changes in *S*_M3_.

**Figure 8.**
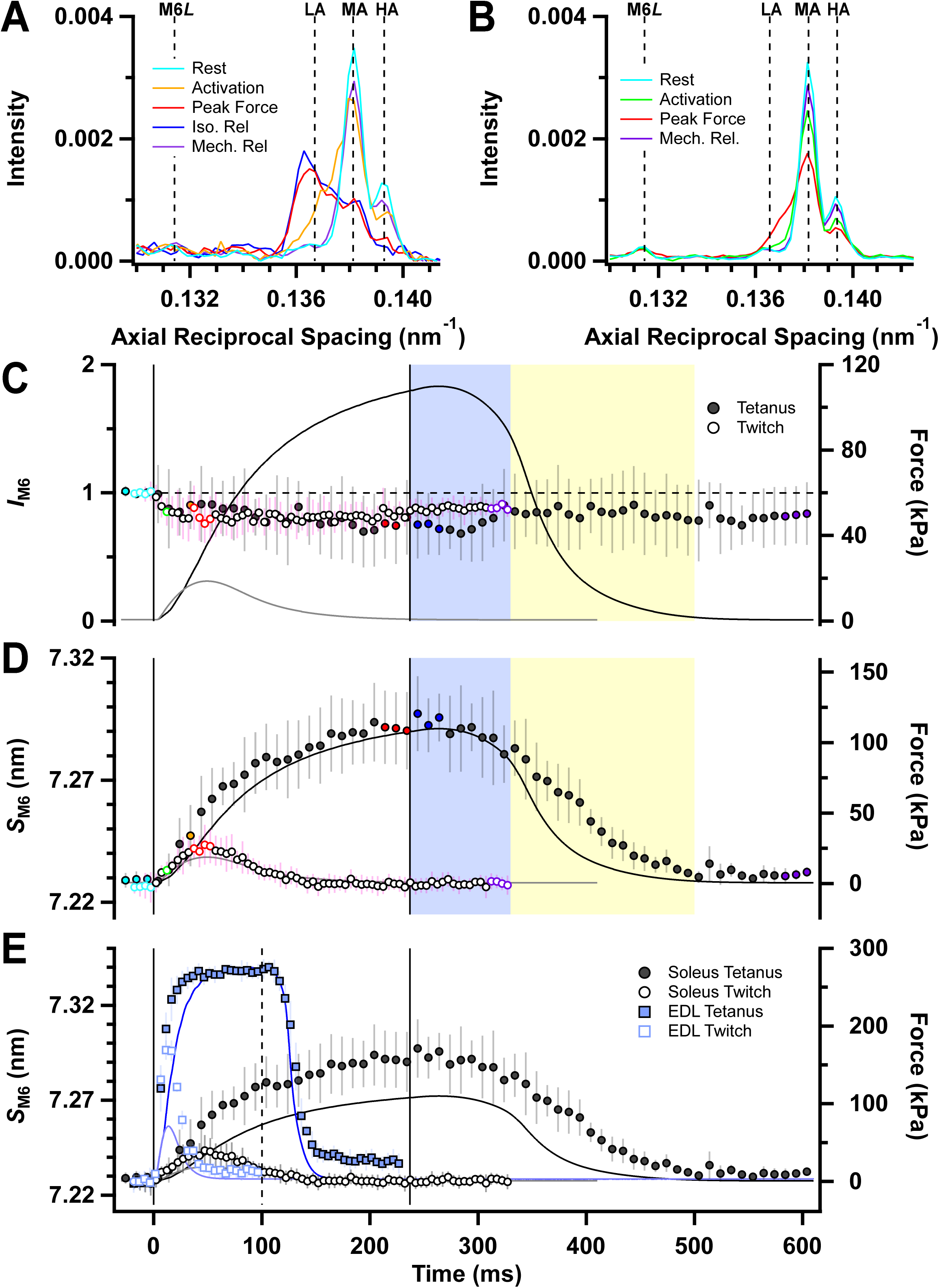
Myosin-based M6 reflection. *A* and *B*, Axial intensity distributions of the M6 reflection in tetanus (*A*) and twitch (*B*). Vertical dashed lines denote low-angle (LA), mid-angle (MA), high-angle (HA) peaks of the M6, and M6*L* reflection. Colour coding denotes periods shown in Fig. 1A. Intensity of the M6 reflection (*I*_M6_, *C*) normalised to its resting value, and spacing of the M6 reflection (*S*_M6_, *D*) for tetanus (filled symbols, mean ± SD, n = 4 muscles) and twitch (open symbols, n = 7 muscles) protocols. In *E*, *S*_M6_ for the soleus (circles) is compared against *S*_M6_ from fast-twitch mouse EDL muscle (blue) during the twitch (open squares, mean ± SD, n = 4 muscles) or 100 ms tetanus (filled squares, n = 5 muscles). *S*_M6_ and force (blue traces) in mouse EDL is from Hill *et al*. (2021), but have been re-calculated using the beamline calibration for those data. Intensities in *A*, *B* and *C* corrected for changes in radial width. Vertical solid lines in *C-E* denote the first and last electrical stimulus for each muscle type. Horizontal dashed line in *C* denote resting *I*_M6_. Colour coding denotes periods shown in Fig. 1A.

Many previous X-ray studies of fast skeletal muscle fibres have interpreted changes in the M3 reflection solely in terms of changes in the conformation of the myosin motors, whereas changes in the M6 reflection were thought to have a large contribution from an additional thick filament component (Reconditi *et al*., 2004; Huxley *et al*., 2006). That interpretation was mainly based on the very different responses of the M3 and M6 reflections to rapid length changes applied during contraction, but has been challenged by high resolution cryo-EM structures of cardiac thick filaments (Tamborrini *et al*., 2023; Dutta *et al*., 2023), which indicate that the M6 reflection, like the M3, comes almost entirely from the myosin motors (Koubassova *et al*., 2025). To explore the implications of that hypothesis for the present results, we calculated the ratio *I*_M6_/*I*_M3_ (Fig. 9), which according to that hypothesis would be related to the distribution of motor mass within each 14.4 nm axial repeat. *I*_M6_/*I*_M3_ increases transiently at the start of a tetanus in rat soleus muscle (Fig. 9A) but returns to close to the resting value at peak force in the tetanus, before increasing again transiently during exponential relaxation. The resting value of *I*_M6_/*I*_M3_ was higher in the batch of muscles used for the twitch experiments, but when the values for twitch and tetanus are normalised to the resting value in each batch (Fig. 9C), the time course of *I*_M6_/*I*_M3_ in twitch and tetanus are remarkably similar. In contrast with the increase in *I*_M6_/*I*_M3_ seen in twitch and tetanus of rat soleus muscle, *I*_M6_/*I*_M3_ *decreases* in a tetanus of mouse EDL muscle (Fig. 9 B,D), although it increases transiently both at the start of activation and during exponential relaxation in a tetanus, and increases in a twitch. The distribution of myosin motor conformations during tetanic contraction of rat soleus muscle is markedly different from that in mouse EDL muscle, as discussed below.

**Figure 9.**
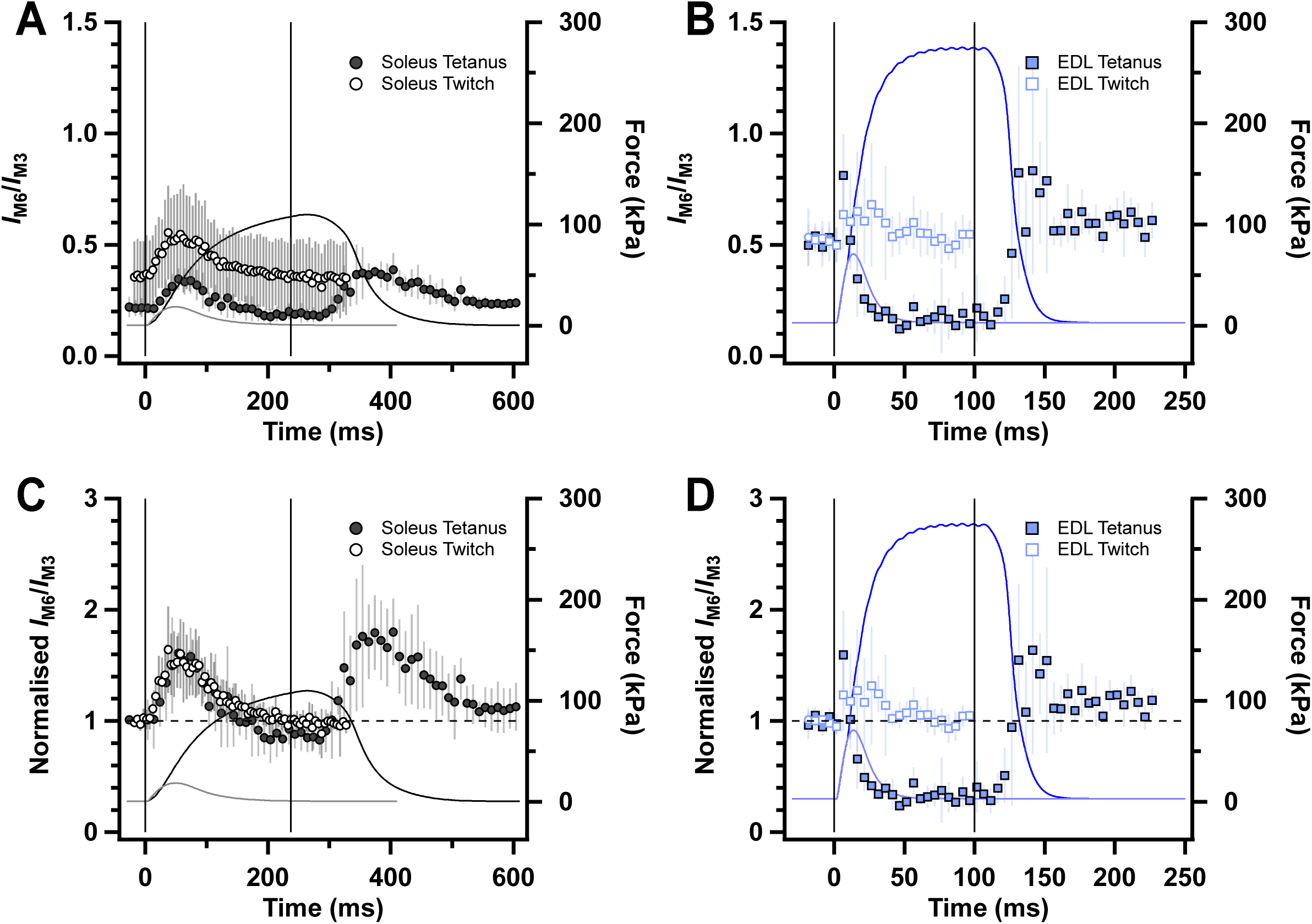
Ratios of the intensities of the M6 and M3 reflections. The intensity of the M6 with respect to the M3 reflection (*I*_M6_/*I*_M3_) for rat soleus (*A* and *C*, grey) and mouse EDL (*B* and *D*, blue, from Hill et al., 2021) during fixed-end tetanus (filled symbols) and twitch (open symbols). Absolute values of *I*_M6_/*I*_M3_ (*A* and *B*) and values normalised with respect to rest (*C* and *D*), superimposed on force. Mean ± SD for n = 7 muscles for soleus twitch (open grey circles), n = 5 muscles for EDL tetanus (filled blue squares) and n = 4 muscles for soleus tetanus (filled grey circles) and EDL twitch (open blue squares). Vertical solid lines in *A*-*D* denote the first and last electrical stimulus. Horizontal dashed lines in *C* and *D* denote resting values.

## Discussion

The main aim of the present experiments was to determine the extent of thick filament activation and the fraction of myosin motors attached to thin filaments during fixed-end contractions of slow skeletal muscle and, by comparing the results with previous studies of fast muscle, to elucidate the molecular structural basis of the lower active force and rate of ATP utilisation in slow muscle. We chose rat soleus muscles contracting at 27°C for these experiments because they contain predominantly type-1 slow fibres, and because the rate of ATP utilisation during fixed-end contraction has been measured at this temperature in both intact and demembranated fibres (Gibbs & Gibson, 1972; Barclay *et al*., 1993; Bottinelli *et al*., 1994). Moreover, the force produced by rat soleus muscles at 27°C is the same as that at physiological temperature, in both twitches and tetani (Close & Hoh, 1968). We used our previous studies of mouse EDL muscles contracting at 27°C (Hill *et al*., 2021, 2022, 2025) to compare results of fast versus slow muscle because those studies used identical methods on a mammalian muscle preparation in conditions in which ATP utilisation and thin filament regulation are also well characterised (Barclay *et al*., 1993; Bottinelli *et al*., 1994; Baylor & Hollingworth, 2003). Apart from differences in kinetics, the changes in the small-angle X-ray diffraction pattern from mouse EDL muscles during fixed-end contraction at 27°C are similar to those in the large number of previous studies on fast muscle fibres of the frog, as discussed in Hill *et al*. (2021).

We used small-angle X-ray diffraction to measure thick filament activation and the attachment of myosin motors to thin filaments in slow muscle. This technique provides molecular structural information about the myosin-containing thick filaments and actin-containing thin filaments in intact contracting muscles on the physiological timescale. Although quantitative interpretation of the observed changes in the X-ray pattern generally depends on molecular structural data from studies of isolated proteins or filaments, the extensive previous studies of the X-ray patterns during contraction of fast muscles from amphibians and mammals provides an established paradigm for the interpretation of the changes observed in slow mammalian muscles, together with its limitations. The multiple X-ray signals available from a single diffraction pattern provides some internal checks.

The increase in the ratio of the intensities of the 11 and 10 reflections (*I*_11_/*I*_10_; Fig. 3C) is often used as a signal for the movement of the myosin motors from the vicinity of the surface of the thick filaments towards the thin filaments during contraction. The increase in *I*_11_/*I*_10_ in a fixed-end tetanus of slow muscle (Fig. 3C) is slightly smaller than that reported previously in mouse EDL muscle, but this cannot be reliably interpreted as a lower fraction of myosin motors attached to thin filaments because *I*_11_/*I*_10_ also depends on other aspects of motor conformation and on the disorder of filament positions within the lattice. The volume of the filament lattice (Fig. 3E), determined from simultaneous measurements of filament lattice spacing and sarcomere length, decreases by about 10% during a tetanus in slow muscle (Fig. 3E), similar to that in mouse EDL muscle (Hill *et al*., 2025), suggesting that similar radial forces are generated in the two muscle types.

The feature of the small-angle X-ray diffraction pattern most directly related to the fraction of myosin motors attached to thin filaments during contraction is the intensity of the first actin layer line (*I*_AL1_), because *I*_AL1_ increases when the myosin catalytic domains attach to actin motors and take up the periodicity of the actin helix (Koubassova *et al*., 2008). Measurements of *I*_AL1_ are complicated by the overlap of AL1 with the nearby first myosin layer line (ML1), but analysis of the mixed layer line at high radius, where myosin contribution is negligible, suggested that *I*_AL1_ increases by only about 40% in a tetanus of rat soleus muscle (Fig. 4C), much less than the more than two-fold increase seen in mouse EDL muscle (Caremani *et al*., 2021; Hill *et al*., 2021). The relatively small increase in *I*_AL1_ observed during a tetanus of rat soleus muscle implies that less than 10% of the myosin motors are attached to thin filaments (Koubassova *et al*., 2008), compared with 25% in fast muscle (Caremani *et al*., 2021; Hill *et al*., 2021). The increase in the intensity of the sixth actin layer line (*I*_AL6_; Fig. 4D) is also much smaller in rat soleus than mouse EDL muscle (Hill *et al*., 2025).

The time course of the change in the intensity of the M3 reflection (*I*_M3_) during a tetanus in rat soleus muscle (Fig. 7C) provides independent support for the conclusion that fewer myosin motors are attached to actin in slow muscle. The initial decreasing phase of *I*_M3_ is associated with the disruption of the folded OFF state of the myosin motors, considered further below, but the later rising phase signals attachment of myosin motors to actin with their long axes roughly perpendicular to the filament axis (Irving *et al*., 2000; Reconditi *et al*., 2011). The rising phase is much smaller in rat soleus muscles than in mouse EDL, in which *I*_M3_ at peak force in the tetanus is more than three times the resting value (Caremani *et al*., 2021; Hill *et al*., 2021). Scaled to their respective resting values, the amplitude of the rising phase in soleus is only about 10% of that in EDL. Because X-ray reflection intensities are proportional to the square of the number of diffractors in a given conformation, the comparison suggests that the fraction of motors attached to actin in slow muscle is only about a third of than in fast muscle, in reasonable agreement with the conclusion from *I*_AL1_.

The most direct estimate of the degree of activation of the thick filaments in muscle is the intensity of the first myosin-based layer line (*I*_ML1_), which signals the helical order of the myosin motors in the folded OFF state (Huxley & Brown, 1967; Linari *et al*., 2015; Irving, 2017). The measurement and even the definition of *I*_ML1_ is complicated in rat soleus muscle because the myosin-based layer lines are sampled by the filament lattice (Squire *et al*., 2004; Ma *et al*., 2019; Koubassova *et al*., 2022). We used two independent approaches to estimate the change in the intensity of the first myosin layer line in the region expected to be dominated by the OFF myosin heads, as described in Results (“*I*_ML1_”, Fig. 4E; *I*_111+201_, Fig. 6B). Both methods suggested that the diffracted intensity associated with OFF motors at the peak of a tetanus in rat soleus muscle is about 50% of that at rest, compared with only 10% in mouse EDL muscle (Hill *et al*., 2021). Since the diffracted intensity is proportional to the square of the number of contributing diffractors, these results imply that only about 30% of the myosin motors are released from the folded OFF conformation at the peak of the tetanus in soleus muscle, compared with about 70% in EDL. Assuming that the fraction of OFF motors is the same in fast and slow muscles at rest, fewer myosin motors are released from the OFF state during a tetanus in slow muscle to become available for binding to thin filaments, active force generation and ATP hydrolysis.

The axial periodicity of the thick filament, signalled by *S*_M3_ (Fig. 7D) and *S*_M6_ (Fig. 8D), has also been widely used as a signal for the degree of activation of the thick filament. *S*_M3_ and *S*_M6_ are about 1.5% larger at the tetanus plateau than at rest in fast muscles from both amphibians (Haselgrove, 1975; Linari *et al*., 2000) and mammals (Hill *et al*., 2021), signalling a change in the packing of the myosin tails in the thick filament backbone on filament activation (Irving, 2017). *S*_M3_ and *S*_M6_ increased by 0.81% and 0.85% respectively at peak force in the tetanus in rat soleus muscle (Table 1; Gong *et al*., 2022), about half of the increases observed in mouse EDL muscle (Fig. 7E, Fig. 8E), and consistent with the conclusion that thick filaments are less fully activated in slow muscle.

The mean peak tetanic force in rat soleus muscle in the conditions of the present experiments was about 110 kPa. Myofibrils occupy about 80% of the cross-sectional area of the muscle (Percario *et al*., 2018), so the myofibrillar force per cross-sectional area is 138 kPa. The cross-sectional area per thick filament, calculated from the dimensions of the filament lattice (Fig. 3D), is 1667 nm^2^, so the force per thick filament is about 230 pN. If 10% of the 294 motors in each half thick filament are attached to actin and generating force as estimated above, the average force per attached motor in isometric conditions is 230/29.4, or about 8 pN, which within the uncertainties of the different techniques is consistent with estimates from single molecule studies (Woody *et al*., 2019; Shchepkin *et al*., 2020). The strain of an actin-attached myosin motor in soleus muscle is 4.5 nm (Caremani *et al*., 2022), so its stiffness is about 1.8 pN.nm^-1^. This value is about three times larger than that estimated by comparing the stiffness of demembranated fibres from slow muscle during active contraction and in rigor (Brenner *et al*., 2012; Percario *et al*., 2018), but consistent with the value obtained by applying the same method to fast muscle. The stiffness of isolated fragments of both fast and slow muscle myosin have also been measured in the optical trap. Studies using fast myosin have consistently reported values larger than 1 pN.nm^-1^ (Lewalle *et al*., 2008), but some studies on myosin from slow muscle reported a lower value, about 0.4 pN.nm^-1^ (Capitanio *et al*., 2006; Brenner *et al*., 2012). A later more comprehensive study suggested that the stiffness of fast and slow myosins is the same in the nucleotide free-state, about 1.5 pN.nm^-1^, but slow myosin is about three times less stiff when ADP is bound (Wang *et al*., 2020), which may explain the different results obtained in the earlier single molecule studies on slow myosins.

In the context of the alternative explanations for the lower force production of slow than fast skeletal muscle summarised in the Introduction, the present therefore results provide strong support for the simpler hypothesis that the force produced by slow muscle is lower because fewer myosin heads are attached to actin filaments, with the corollary that the force and stiffness of an attached myosin head are the same for the two myosin isoforms. That conclusion is also consistent with the energetics of muscle shortening against a load. Soleus muscles can convert the chemical energy of ATP hydrolysis into mechanical work with an efficiency of about 50% at the temperature of the present experiments (Barclay *et al*., 2010*a*). The working stroke of isolated slow myosin motors is about 6 nm, not significantly different from that of fast type-2 myosins (Capitanio *et al*., 2006). With a force per motor of 8 pN, as estimated from the present experiments, each motor would therefore produce 48 pN·nm of work during a single interaction with actin, which is 48% of the 100 pN.nm available from the hydrolysis of one molecule of ATP in mammalian muscle (Barclay *et al*., 2010*b*), in agreement with the measured mechanical efficiency of slow muscle (Barclay *et al*., 2010*a*), and with similar estimates for fast muscle (Piazzesi *et al*., 2007; Barclay *et al*., 2010*a*).

The slower rate of active force development in slow muscle following tetanic stimulation allows a clear temporal separation between fast X-ray signals associated with thin filament activation and the slower signals associated with force development. The latter include the ratio of the intensities of the 11 and 10 equatorial reflections (*I*_11_/*I*_10_; Fig. 3C), the rising phase of the change in the intensity of the M3 reflection (*I*_M3_; Fig. 7C), the increase in thick filament periodicity associated with thick filament activation signalled by *S*_M3_ (Fig. 7D) and *S*_M6_ (Fig. 8D), and the intensity of the first myosin layer line associated with the folded OFF conformation of the myosin motors (“*I*_ML1_”, Fig. 4E; *I*_111+201_, Fig. 6B). All these signals are greatly attenuated in a twitch compared with those in a tetanus. Given the six-fold lower peak force in the twitch, this result is consistent with previous X-ray studies of fast skeletal muscle that established an association of *I*_11_/*I*_10_ and the rising phase of *I*_M3_ with the number of actin-attached force-generating motors, and that of *S*_M3_, *S*_M6_ and *I*_ML1_ with thick filament stress.

The fast signals include the early decrease in the intensities of the M3 reflection (*I*_M3_; Fig. 7C), the myosin-based reflections *I*_103_ (Fig. 6E) and *I*_113+203_ (Fig. 6F) and the M6 reflection (*I*_M6_; Fig. 8C), and the increase in *I*_AL1_ (Fig. 4C). The decrease in *I*_M3_ has a half time of about 20 ms in a tetanus of rat soleus muscle (Table 2), compared with 72 ms for force development. The other fast signals are synchronous with the *I*_M3_ decrease within the resolution of the measurements. Moreover, all these signals have the same amplitude in a twitch and a tetanus, as expected from the finding that thin filament activation is complete, albeit transiently, in the twitch (Baylor & Hollingworth, 2003) but in marked contrast with the six-fold higher peak force in the tetanus. The fast decreases in *I*_M3_, *I*_M6_, *I*_103_, *I*_M3_, and *I*_113+203_, which are associated with the axial mass distribution of the myosin motors, show that thin filament activation is accompanied by a fast change in the conformation of the myosin motors that precedes actin attachment and force generation, implying the existence of a fast signalling pathway between the thin and thick filaments that is distinct from the mechano-sensing mechanism of myosin filament activation that operates in fast muscle (Linari *et al*., 2015; Irving, 2017; Hill *et al*., 2022).

This inter-filament signalling pathway might be mediated by ‘constitutively ON’ or ‘sentinel’ myosin motors (Linari *et al*., 2015; Irving, 2017; Craig & Padrón, 2022), which are immediately available for actin binding on thin filament activation. Another possibility is mediation by myosin binding protein-C (MyBP-C), which is composed of C-terminal domains that are an integral part of the thick filament backbone linked to N-terminal domains that may bind to thin filaments in resting muscle (Luther *et al*., 2011; Tamborrini *et al*., 2023; Dutta *et al*., 2023). The N-terminal domains of MyBP-C might detach from the thin filaments on activation, triggered either by thin filament activation itself or by the roughly 25 nm filament sliding which occurs even in a fixed-end twitch (Fig. 2C) as a result of tendon compliance. Further small-angle X-ray diffraction experiments involving mechanical perturbations will be required to test these hypotheses.

In summary, the X-ray diffraction data presented here show that the molecular structural changes associated with twitch and tetanic contractions of rat soleus muscles are not simply a slower version of those previously described in fast muscles. Fewer myosin motors are activated in a tetanus in slow muscle, and fewer motors bind to actin. The structural changes in the thin and thick filaments of soleus muscles on activation can be temporally resolved into two classes, fast changes triggered by thin filament activation and slow changes associated with active force generation and thick filament stress.

## Acknowledgements

We would like to thank the ID02 technical staff, Narayanan Theyencheri, Laurent Jacqmin, Peter Boesecke and Michael Sztucki, and the ID17 Biomedical Facility staff, Mélanie Jomard, Andrea Tramond and Michael Kirsch (European Synchrotron Radiation Facility), for their support during the beamtime; the European Synchrotron Radiation Facility for the award of synchrotron beamtime; the I22 technical staff, Olga Shebanova, Tim Snow, and Nick Terrill (Diamond Light Source), for their support during the beamtime; the Diamond Light Source for the award of synchrotron time; Kevin Whitehill (Public Health England) for his assistance with animal care; Kawal Rhode and Zhouyang Xu (King’s College London) for their assistance with the 3D printing of the muscle trough. This work was funded by the Medical Research Council (MR/R01700X/1), the European Synchrotron Radiation Facility and the Diamond Light Source. M.K. was supported by the Wellcome Trust (215482/Z/19/Z) award to M.I. A.A. was supported by a British Heart Foundation PhD fellowship (FS/4yPhD/F/21/34154) and King’s British Heart Foundation Centre of Research Excellence Award RE/18/2/34213. Y.W. and E.B. were supported by a British Heart Foundation Intermediate Basic Science Research Fellowship awarded to E.B. (FS/17/3/32604). L.F. was funded by a Sir Henry Dale Fellowship awarded by the Wellcome Trust and the Royal Society (210464/Z/18/Z).

---

**Figure S1.**
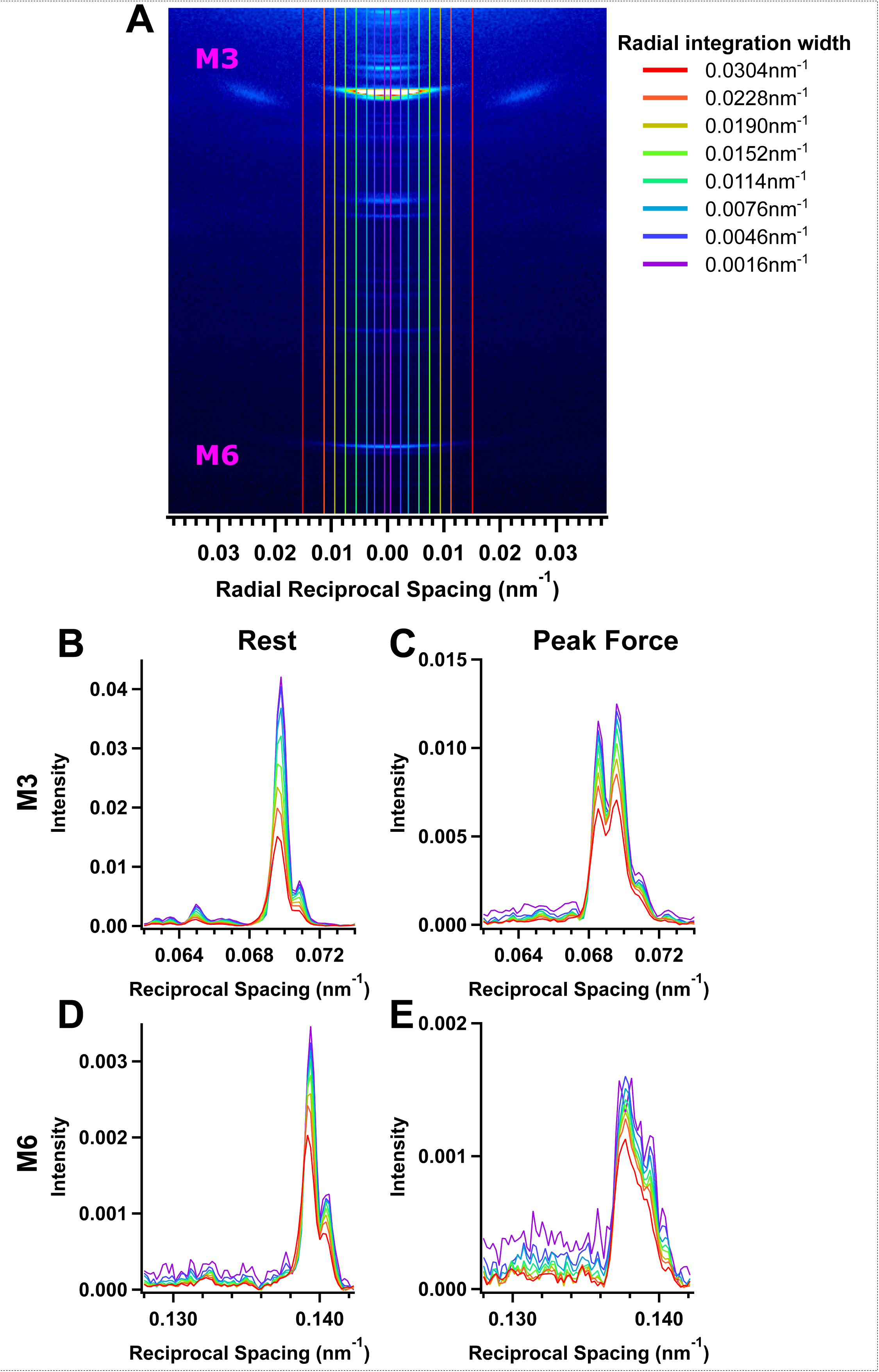
Sarcomere reflections. Ultra-small angle axial intensity distributions, plotted as measured intensity times the square of reciprocal spacing, from resting muscle with sample-to-detector distance of 31 m at beamline ID02 at ESRF (black) and 8.2 m at beamline I22 at Diamond Light Source (red). Vertical dashed black indicates the orders of the sarcomere reflections. Average data from n = 7 muscles (black) and n = 4 muscles (red).

**Figure S2.**
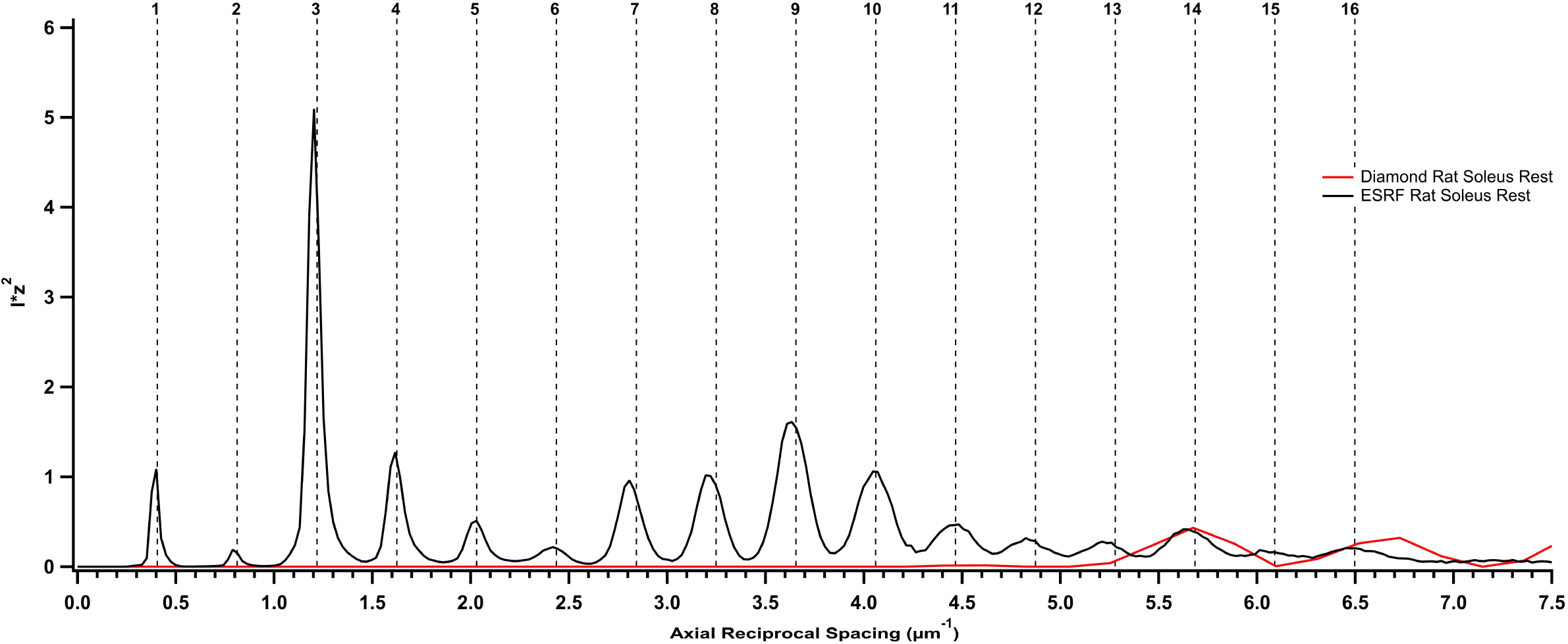
Optimisation of radial integration limits for the meridional reflections. *A*, region of the two-dimensional small-angle X-ray diffraction in Fig. 1E containing the M3 and M6 reflections (magenta labels). Vertical lines denote different radial integration limits. *B-E*, Axial intensity distributions using the different radial limits for the M3 (*B* and *C*) and M6 (*D* and *E*) reflections at rest (*B* and *D*) and peak tetanic force (*C* and *E*). Profiles corrected by the cross-meridional width (see *Methods*). Average data from of n = 4 muscles.

**Figure S3.**
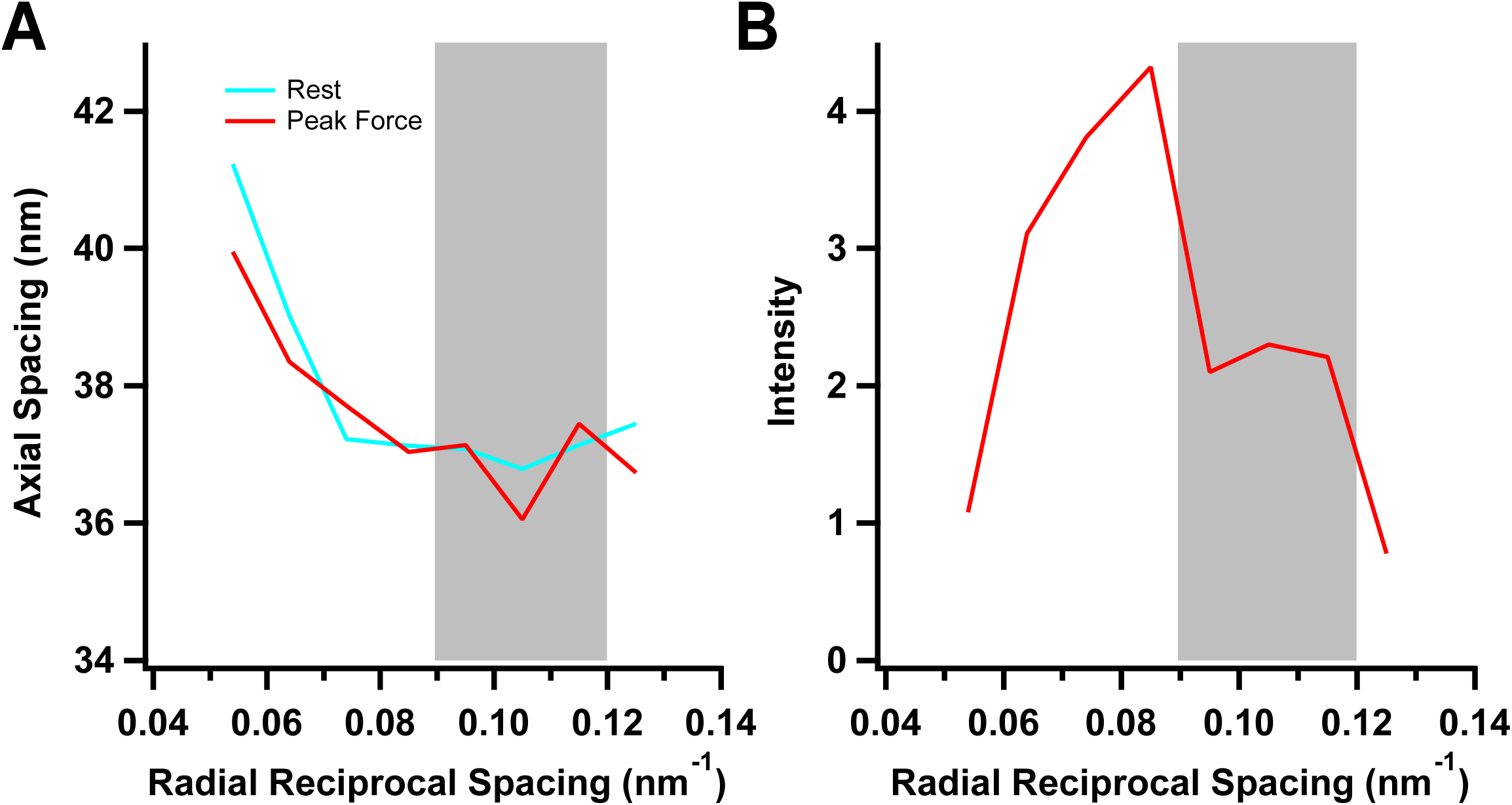
Axial spacing and intensity of the mixed first-order actin and myosin layer line reflection in the fixed-end tetanus. Spacing (*A)* and normalised intensity (*B*) of the mixed first order actin and myosin layer line reflections at rest (cyan) and peak force (red). Grey shaded region in *A* and *B* corresponds to the radial integration limits used to obtain *I*_AL1_ in Fig. 4D. See *Methods* for detailed overview.

**Table S1.** Beamline properties and detector specifications.

| Property | European Synchrotron<br>Radiation Facility; ID02 | Diamond Light Source; I22 |
| --- | --- | --- |
| Flux (photons.s<sup>-1</sup>) | ~2 x 10<sup>13</sup> | ~6 x 10<sup>12</sup> |
| Wavelength (nm) | 0.1 | 0.1 |
| FWHM (μm; h x v) | ~230 x ~50 | ~300 x ~100 |
| Detector model, manufacturer | Eiger 2X 4M, Dectris | Pilatus P3 2M, Dectris |
| Modules (h x v) | 2 x 4 | 3 x 8 |
| Intermodule gap (px; h, v) | 12, 38 | 7, 17 |
| Active area (mm; h x v) | 155.1 x 162.15 | 253.7 x 288.8 |
| Pixel array (px; h x v) | 2068 x 2162 | 1475 x 1679 |
| Pixel size (μm; h x v) | 75 x 75 | 172 x 172 |

**Table S2.** Tetanus ANOVA and post-hoc test p-values.

| Parameter | Main Effect | Rest vs |  |  | Peak Force vs |  | Isometric Relaxation vs |
| --- | --- | --- | --- | --- | --- | --- | --- |
|  |  | Peak Force | Isometric Relaxation | Mechanically Relaxed | Isometric Relaxation | Mechanically Relaxed | Mechanically Relaxed |
| Force (kPa) | <0.001 | 0.009 | 0.008 | 0.733 | 0.167 | 0.009 | 0.008 |
| $d_{10}$ (nm) | <0.001 | 0.047 | 0.075 | <0.001 | <0.001 | 0.551 | 0.333 |
| $I_{ML1}$ | <0.001 | <0.001 | <0.001 | 0.007 | 0.297 | 0.005 | <0.001 |
| $A_{ML1}$ | <0.001 | <0.001 | <0.001 | 0.010 | 0.749 | 0.005 | <0.001 |
| $I_{AL1}$ | 0.009 | 0.164 | 0.064 | 0.781 | 0.815 | 0.362 | 0.197 |
| $I_{AL6}$ | <0.001 | <0.001 | <0.001 | 0.785 | 0.572 | 0.004 | <0.001 |
| $I_{101}$ | <0.001 | 0.004 | <0.001 | 0.108 | 1.000 | 0.068 | 0.010 |
| $I_{111+201}$ | <0.001 | <0.001 | <0.001 | 0.014 | 1.000 | <0.001 | <0.001 |
| $I_{102}$ | <0.001 | <0.001 | <0.001 | 0.007 | 0.998 | 0.152 | 0.133 |
| $I_{112+202}$ | <0.001 | <0.001 | <0.001 | 0.001 | 1.000 | <0.001 | 0.017 |
| $I_{103}$ | <0.001 | <0.001 | <0.001 | <0.001 | 1.000 | 0.009 | 0.031 |
| $I_{113+203}$ | <0.001 | <0.001 | <0.001 | 0.002 | 1.000 | 0.774 | 1.000 |
| $I_{M6}$ | <0.001 | 0.048 | 0.009 | 0.067 | 0.692 | 0.236 | 0.017 |
| $S_{M6}$ (nm) | <0.001 | <0.001 | <0.001 | 0.139 | 0.329 | <0.001 | <0.001 |
| $I_{M3}$ | 0.001 | 0.274 | 0.178 | <0.001 | 0.671 | 0.120 | 0.112 |
| $A_{M3}$ | <0.001 | 0.207 | 0.135 | <0.001 | 0.768 | 0.132 | 0.118 |
| $S_{M3}$ (nm) | <0.001 | <0.001 | <0.001 | 0.101 | 0.233 | <0.001 | <0.001 |
| $I_{11}/I_{10}$ | <0.001 | <0.001 | <0.001 | 0.044 | 0.557 | <0.001 | <0.001 |

**Table S3.** Tetanus ANOVA and post-hoc test p-values.

| Parameter | Main Effect | Rest vs |  | Peak Force vs |
| --- | --- | --- | --- | --- |
|  |  | Peak Force | Mechanically Relaxed | Mechanically Relaxed |
| Force (kPa) | <0.001 | <0.001 | 0.156 | 0.161 |
| $d_{10}$ (nm) | 0.025 | 0.145 | <0.001 | 0.126 |
| $I_{ML1}$ | 0.004 | 0.017 | 0.112 | 0.158 |
| $A_{ML1}$ | 0.002 | 0.012 | 0.101 | 0.120 |
| $I_{AL1}$ | <0.001 | <0.001 | 0.043 | <0.001 |
| $I_{AL6}$ | <0.001 | 0.016 | 0.284 | 0.007 |
| $I_{101}$ | <0.001 | <0.001 | 0.022 | <0.001 |
| $I_{111+201}$ | <0.001 | <0.001 | 0.015 | <0.001 |
| $I_{102}$ | <0.001 | <0.001 | 0.052 | 0.002 |
| $I_{112+202}$ | <0.001 | <0.001 | 0.048 | <0.001 |
| $I_{103}$ | <0.001 | <0.001 | 0.113 | <0.001 |
| $I_{113+203}$ | <0.001 | <0.001 | 0.679 | <0.001 |
| $I_{M6}$ | <0.001 | <0.001 | <0.001 | 0.017 |
| $S_{M6}$ (nm) | <0.001 | <0.001 | 0.031 | <0.001 |
| $I_{M3}$ | <0.001 | <0.001 | <0.001 | <0.001 |
| $A_{M3}$ | <0.001 | <0.001 | <0.001 | <0.001 |
| $S_{M3}$ (nm) | 0.007 | 0.009 | 0.002 | 0.070 |
| $I_{11}/I_{10}$ | <0.001 | <0.001 | 0.002 | <0.001 |

**Table S4.**
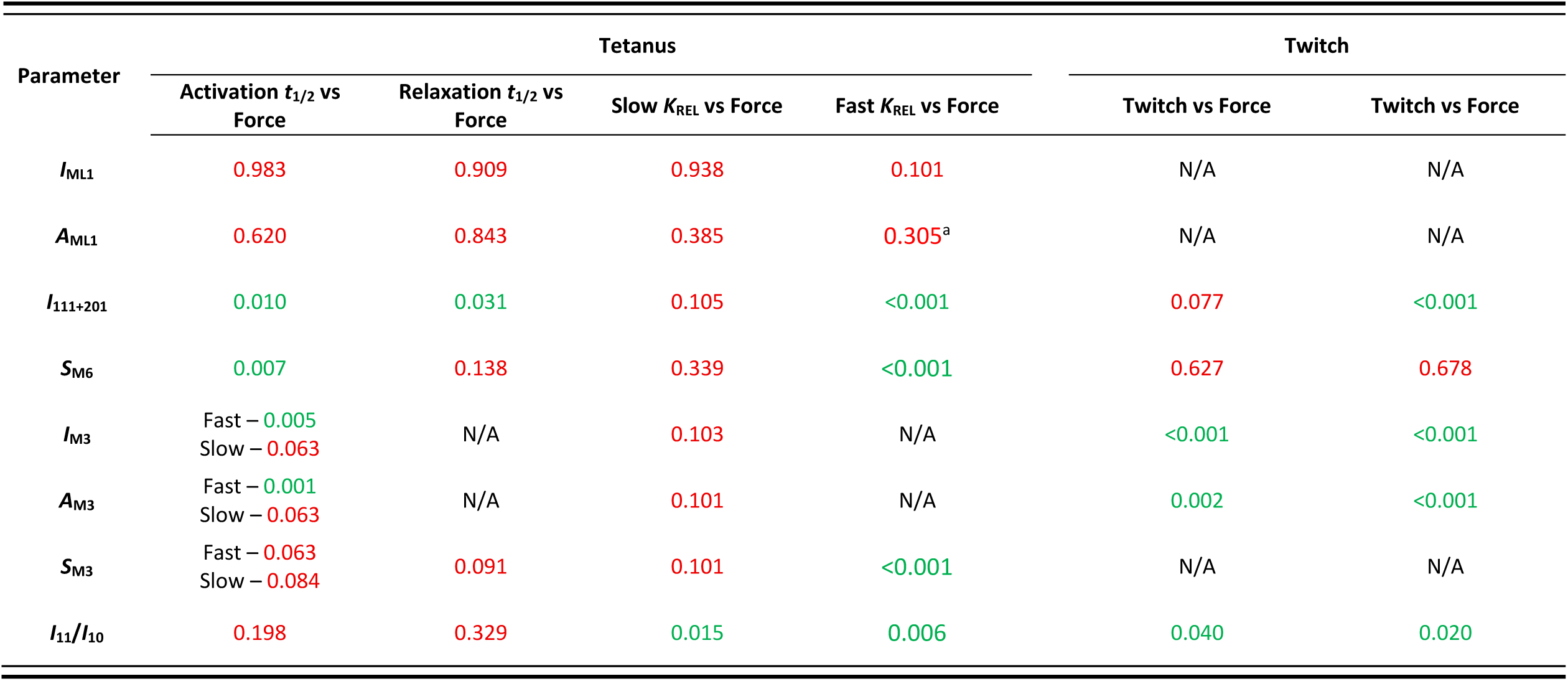
Activation and relaxation half time and rate constant *t*-test p-values.

| Parameter | Tetanus |  |  |  | Twitch |  |
| --- | --- | --- | --- | --- | --- | --- |
| | Activation $t_{1/2}$ vs Force | Relaxation $t_{1/2}$ vs Force | Slow $K_{REL}$ vs Force | Fast $K_{REL}$ vs Force | Twitch vs Force | Twitch vs Force |
| $I_{ML1}$ | 0.983 | 0.909 | 0.938 | 0.101 | N/A | N/A |
| $A_{ML1}$ | 0.620 | 0.843 | 0.385 | 0.305<sup>a</sup> | N/A | N/A |
| $I_{111+201}$ | 0.010 | 0.031 | 0.105 | <0.001 | 0.077 | <0.001 |
| $S_{M6}$ | 0.007 | 0.138 | 0.339 | <0.001 | 0.627 | 0.678 |
| $I_{M3}$ | Fast – 0.005<br>Slow – 0.063 | N/A | 0.103 | N/A | <0.001 | <0.001 |
| $A_{M3}$ | Fast – 0.001<br>Slow – 0.063 | N/A | 0.101 | N/A | 0.002 | <0.001 |
| $S_{M3}$ | Fast – 0.063<br>Slow – 0.084 | 0.091 | 0.101 | <0.001 | N/A | N/A |
| $I_{11}/I_{10}$ | 0.198 | 0.329 | 0.015 | 0.006 | 0.040 | 0.020 |

